# Combining DNA scaffolds and acoustic force spectroscopy to characterize individual protein bonds

**DOI:** 10.1101/2022.08.14.503897

**Authors:** Yong Jian Wang, Claire Valotteau, Adrien Aimard, Lorenzo Villanueva, Dorota Kostrz, Maryne Follenfant, Terence Strick, Patrick Chames, Felix Rico, Charlie Gosse, Laurent Limozin

## Abstract

Single-molecule data are of great significance in biology, chemistry, and medicine. However, experimental tools to characterize, in a multiplexed manner, protein bond rupture under force are needed. Acoustic force spectroscopy (AFS) is an emerging manipulation technique which generates acoustic waves to apply force in parallel on a large population of microbeads tethered to a surface. We have exploited this configuration on a recently developed modular Junctured-DNA (J-DNA) scaffold designed to study protein-protein interactions at the single-molecule level. By applying repetitive constant force steps on the FKBP12-rapamycin-FRB complex, we measured its unbinding kinetics under force at the single-bond level. Special effort was made in analyzing the data in order to identify potential pitfalls. We established a calibration method allowing in situ force determination during the course of the unbinding measurement. We compare our results with well established techniques, such as magnetic tweezers, to ensure their accuracy. We also apply our strategy for measuring the force dependent rupture of a single domain antibody with its antigen. We get a good agreement with standard measurement at zero force. Our technique offers single molecule precision for multiplexed measurements of interactions of biotechnological and medical interest.

## Introduction

Biomolecules binding properties are governing biological phenomena and consequently are relevant for evaluating therapeutics. While bulk measurements performed on populations of molecules remain the standard characterization techniques, single-molecule force spectroscopy (SMFS) on individual pairs of interacting partners has emerged as a powerful complementary strategy, because it uniquely gives access to individual bond response to force. This ability is particularly significant for the study of biological phenomena where mechanical forces intervene in bond function, like in hematology [1], neuroscience [2], immunology [3], or cell biology [4]. More precisely, the various SMFS tools [5] can be separated in two categories. On one hand, several techniques scrutinize a single bond at a time, the interacting molecules being mounted in series with a spring-like dynamometer such as in atomic force microscopy (AFM) [6, 7], optical tweezers (OT) [8, 2], or micropipette-based biomembrane force probe [9]. Typically, a force ramp is applied to the bond and the force is determined at the rupture from the extension of the calibrated spring. The dependence of the off-rate as a function of the force is then deduced by a mathematical transformation of the rupture force distributions obtained at different loading rates [10]. Offering a high precision, these methods however suffer from a limited throughput. On the other hand, measurements on several bonds at a time usually involve a spatially extended force field acting on microbeads, a configuration encountered in magnetic tweezers (MT) [11], laminar flow chamber (LFC) [12, 13], centrifugal force microscopy (CFM) [14, 15], dielectrophoretic force spectroscopy [16] and more recently acoustic force spectroscopy (AFS) [17, 18]. In those techniques, the distribution of bond lifetimes can be directly measured at constant force but the range of applied force is often limited. Also, calibration of the force usually relies on a physical model and/or assumes the homogeneity of the force field and bead properties [19].

Whether investigating bonds one by one or in parallel, the stochastic nature of bond rupture requires the accumulation of a large number of events in order to characterize the unbinding distribution. This intrinsic constraint of single-molecule approaches is often experimentally problematic because protein bond characterization suffer from irreversible unbinding: the molecular partners are taken apart by force, preventing any rebinding and therefore limiting the accessible statistics. To address this issue, molecular scaffolds that include a leash have been engineered to keep the reactants in close proximity despite the dissociation, hence favouring rebinding. An additional advantage provided by such constructs is mechanical fingerprinting: upon bond rupture, the two partners must end separated by the length of the leash; events that do not verify this criterion can be easily sorted out and discarded [1, 14, 20, 21]. In practice, these leashes may consist in peptidic chains [1, 2, 4, 22, 23], single DNA strands [24, 25, 26], or double DNA strands, with [15, 27, 28] or without nicks [29, 30, 31]. While peptidic linkers require to design and produce a specific fusion protein, nucleic acids based structures offer more modularity when combined with state-of-the-art DNA-protein coupling strategies [20, 30, 32, 33, 34, 35, 36, 37]. As a matter of fact, the latter kind of scaffold have already been used for non-covalent bond characterization on various SMFS setups: OT [26, 24, 25, 15, 14], MT [30, 25, 29, 31], CFM [15], and LFC [28].

Here, we introduce the combination of a modular DNA scaffold, namely junctured-DNA (J-DNA) [38, 30], with AFS, an emerging parallel method which potentially offers a high dynamic range in force application [17, 18] (Fig. 1). The probed protein complex is attached to both the surface of the flow cell and a bead via kilobase pair-long DNA shanks (Fig. S1); it thus provides a natural way to calibrate the force applied to each pair of partners independently. Indeed one can rely on the inverse pendulum method usually applied in MT [39, 19], with the notable advantage of a direct and accurate determination of the tether length, which is not always possible with MT. Indeed, contrarily to MT, AFS does not impose the orientation of the bead upon pulling, which permits to determine the point of anchoring the DNA on the bead. Additionally, while individual fit of calibration parameters (diffusion and stiffness) can be hampered by the acquisition conditions (camera frame rate) for higher forces, a global fit applied for all acoustic powers for each bead allows to overcome this issue. Thus, simultaneous calibration and unbinding measurements at low camera frame rate are achieved for all forces. This ability to measure the applied force during unbinding experiment is critical because the acoustic field exhibits high variability and spatial heterogeneity [18, 40, 41].

**Figure 1:**
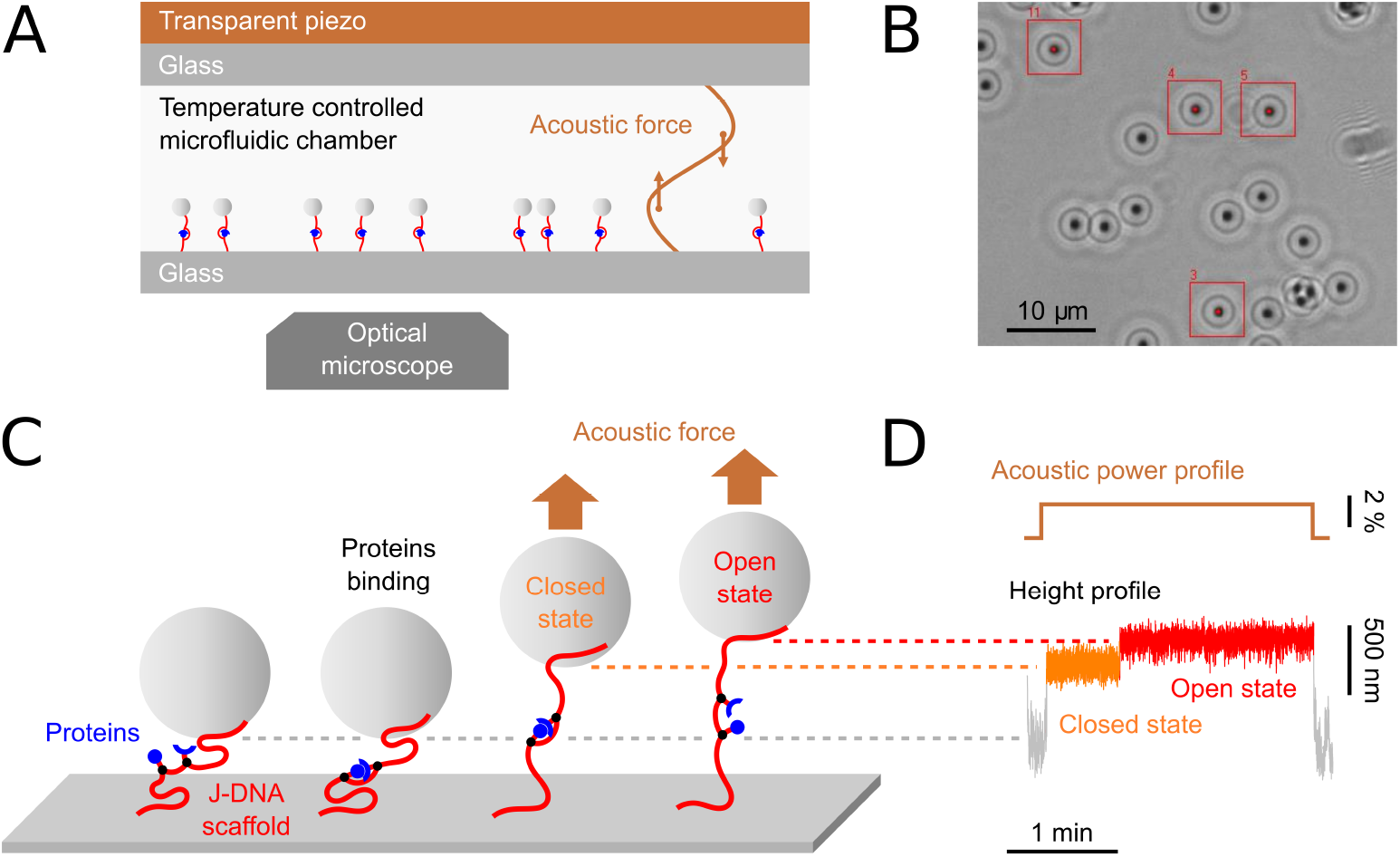
Principle of the method and sample trace. (A) Beads are tethered in a microfluidic chamber equipped with a piezo element, generating acoustic waves, and temperature control (Not to scale). (B) Part of the field of view, showing beads that are selected (red squares) to be tracked in real-time and 3D thanks to their diffraction fringes. (C) The beads are tethered to the chamber surface through a J-DNA (red) scaffold maintaining the protein partners (blue) via a leash. Bond formation occurs in absence of significant acoustic force. Upon application of acoustic force, the extension of the scaffold in closed (proteins bound, orange extension) and open state (proteins unbound, red extension) allows to distinguish the unbinding of the protein complex. (D) The corresponding acoustic power and bead height profiles of the bead.

We first apply our method to the FKBP12-rapamycin-FRB complex, a well-known system that has been studied using numerous biochemical [42, 43, 44] and biophysical [45, 46, 47, 48, 49, 50] techniques, including SMFS [30, 22]. In addition to its interest for benchmarking, rapamycin has an important biomedical relevance, being one of the earliest modulator of protein-protein interaction to have reached to the clinic [51, 52]. Our results successfully compare with the unbinding data obtained by surface plasmon resonance (SPR) [45, 46] and by SMFS [30, 22]. In this latter case, we extend the range of the forces previously explored with MT [30, 22]. Furthermore, the determination of the off-rate vs force dependence for different individual pairs of interacting molecules provides a check of the method accuracy, and a way to look for bond heterogeneity. For a second proof-of-concept, we turn to a model antigen-antibody bond involving a nanobody. Because of their remarkable binding properties [53], those 12kDa antibodies serve as building blocks for bispecific constructs used for cancer immunotherapy [54]. In this context, the bond they form are subjected to mechanical forces [13], so it is important to characterize it under these conditions. An extended force range was critical to achieve off-rate determination of strong bond like nanobody-antigen in a reasonable time. Doing so, the off-rate extrapolated at zero force is found in good agreement with independent determination relying on SPR [55].

## Materials and Methods

### Molecules and reagents

#### Proteins

All proteins were labeled with a single azido group so as to enable their conjugation via click chemistry to 5’ DBCO-modified oligonucleotides, the latter 9-base pair long sequences being used for further site-specific ligation to the J-DNA scaffolds [30]. More precisely, for the study of the rapamycin complex, a 4-azido-L-phenylalanine residue was introduced at the N-terminus of both FKBP12 and FRB proteins by unnatural amino acid incorporation [56, 57, 58], which yielded the previously characterized FKBP12_M0AzF_ and FRB_A2020AzF_ mutants [30]. For the nanobody-antigen study, SdAb19 (later noted Nef19), a nanobody directed against the HIV-1 Nef protein [55] and C-terminally fused to both a c-myc tag and a hexahis one was produced in the periplasm of *E. coli* and purified by immobilized metal affinity purification as previously described [59]. In parallel, a truncated portion (57-205) of HIV-1 Nef was produced in the cytoplasm of *E. Coli* and purified as above. Microbial Transglutaminase (Zedira GmbH, Darmstadt, Germany) and its substrate azido-PEG_3_-amine (click chemistry tools) was used to site specifically add an azide function on the glutamine residue of the c-myc tag of both proteins [60, 61].

#### J-DNA scaffold

Similarly to the others nucleic acids-based scaffolds recently developed for single-molecule investigations [38, 30, 15, 27, 28, 29, 31, 26, 25, 24], J-DNAs have a forceps-like anatomy [20] (Fig. S1A). The interacting protein partners are engrafted at the tips of the nanotool and the handles allowing the manipulation are attached to the extremities of the shanks. The leash, which prevents the whole assembly to be torn apart upon unbinding, connects the two tip-shank segments at the junctions. We here relied on our previous protocol [30] to synthesize two versions of the J-DNAs and functionalized them with the proteins of interest that had been tagged with single-azido group [30]. More specifically, in the present study we used both the asymmetric scaffold described in Kostrz et al. [30] and a new symmetric one (Table S1). In the latter construct the length of the shanks was modified so as to make it easier to discriminate protein-protein interactions from protein-surface ones, i.e. to have extension jumps of different amplitudes for specific binding and for non-specific sticking to the bead or to the chamber (Table S2). Transforming the asymmetric scaffold into the symmetric one was simply achieved by shifting the positions on the template of the primers used in the first PCR reaction of the J-DNA fabrication process (see the caption of Table S1 for the sequences).

#### Beads

Streptavidin-coated silica beads of 1.58 *μ*m diameter (1.0% w/v, Spherotech, Lake forest, IL, USA, cat. SVSIP-15-5) were used for the FKBP12-rapamycin-FRB measurement. For the Nef-Nef19 one, we additionally relied on 3 *μ*m diameter beads (1.0% w/v, Spherotech, cat. SVSIP-30-5).

#### Buffers

Unless otherwise specified, all chemicals for buffer preparation were from Sigma-Aldrich (Merck KGaA, Darmstadt, Germany). Functionalization buffer was prepared using phosphate buffer saline (PBS) tabs, bovine serum albumin (BSA, cat. A7030) and Pluronic F-127 (cat. 9003-11-6). Single use aliquots of rapamycin (cat. 53123-88-9) were prepared by dissolution of the powder in dimethylsulfoxide (DMSO) to a concentration of 50 *μ*M and stored at -80°C. Importantly, due to its instability, this compound was always added extemporaneously. All the solutions were supplemented with 5 mM sodium azide and 0.5 mM ethylenediaminetetraacetic acid (EDTA), filtered at 0.2 *μ*m and stored at 4°C. Assays with J-DNA functionalized with FKBP12 and FRB were conducted in mTOR buffer: 20 mM potassium HEPES, pH 7.8, 100 mM KCl, 5 mM MgCl_2_, 0.1% Tween-20, 0.5 mg/ml BSA, and 2 mM dithiothreitol (DTT), supplemented with 500 nM rapamycin. Assays with J-DNA functionalized with Nef and Nef19 were conducted in PBS, 0.1% Tween-20, and 0.5 mg/ml BSA.

### Preparation of the AFS chamber

The AFS chamber (Lumicks B.V., Amsterdam, Netherlands) consists of two parallel glass slides delineating a 100 *μ*m high fluid channel and approximately 10×2 mm^2^ surface. The internal chmaber volume with tubbing is about 20 *μ*L. A transparent piezoelectric element is present on top of the assembly to generate the acoustic waves that will pull on the beads (Fig. 1A). A syringe connected to the outlet port allows one to suck solutions placed in the inlet reservoir, either manually or with a syringe pump (Aladdin 1000; WPI, Sarasota, Florida, USA).

The AFS chamber surface was silanized to realize a highly hydrophobic surface in order to facilitate further binding of antibodies. To this end, the AFS chamber was first incubated with a piranha solution (2/3 sulfuric acid + 1/3 hydrogen peroxide 50% in water) [12] for 10 minutes, rinsed with water, dried with nitrogen, incubated with Sigmacote® (Sigma-Aldrich, cat. SL2) for 1 min, dried, rinsed with acetone (Sigma) and rinsed with PBS. The AFS chamber was next functionalized with polyclonal anti-digoxigenin from sheep (Sigma, Roche cat. 11333089001) by injection of 50 *μ*L of a 200 *μ*g/mL anti-Dig solution (20 min incubation). The AFS chamber was finally double-passivated by flushing first 200 *μ*L of 0.2% PBS-BSA (30 min incubation), and then 200 *μ*L of 0.5% PBS-Pluronic solution (30 min incubation).

For the measurements on the FKBP12-rapamycin-FRB complex, the AFS chamber was rinsed with 200 *μ*L of mTOR buffer; then the J-DNA on which the proteins had been pre-assembled were injected at 0.5-1pM concentration in mTOR buffer (30 min incubation). Anchoring of the scaffold to the anti-digoxigenin glass surface is mediated by digoxigenin molecules located at one extremity of one of the shanks (Fig. S1A). Streptavidin-coated beads were washed two times by spinning down 10 *μ*L of solution at 0.5 % w/v diluted in 1 mL of mTOR buffer. After a final resuspension in 50 *μ*L of mTOR supplemented with 500 nM of rapamycin, the particles were injected in the AFS chamber and allow to settle for few minutes so that they can bind to the extremity of the second shank, which is decorated with biotin molecules (Fig. S1A). A gentle flow of rapamycin containg mTOR buffer was then applied to remove the untethered beads.

For Nef-Nef19 measurements, the AFS chamber was only passivated by injection of 0.2 mL of 0.2% BSA solution in PBS for 60 minutes incubation. The prefunctionalized J-DNA diluted at 1pM in the same buffer was then injected in the AFS chamber (30 min incubation), before another rinsing with 0.2 mL PBS-BSA. Beads were washed three times by spinning down 50 *μ*L of solution at 1% w/v diluted in 1 mL of 0.2% BSA solution in PBS. After final resuspension in 50 *μ*L of 0.2% BSA in PBS, they were injected in the AFS chamber and incubated for few minutes. A gentle flow of 0.2% BSA in PBS was finally applied to remove the untethered beads.

### Microscope and AFS setup

The sample is illuminated by a fiber-coupled LED (M660F1, Thorlabs, Newton, NJ, USA) and imaged using an inverted microscope (IX71, Olympus, Rungis, France) equipped with a 20× air objective (Uplan F, Olympus) and an USB-camera (UI1324, IDS, Obersulm, Germany) capable of acquiring 1936 × 1216 frames at 55 Hz which corresponds to a 458 *μ*m x 420 *μ*m field of view. Higher frame rates can be achieved for smaller areas. The AFS chamber is screwed on a sample holder, which vertical position can be adjusted by a nanometer step motor (M-110.12S, PI, Karlsruhe, Germany) driven by a digital controller.

The piezo of the AFS chip is driven by a function generator (Lumicks B.V., generation 3) controlled by a computer (via USB) and a Labview (National Instruments, Texas) interface. The application to the piezo of an alternative tension in the MHz frequency range creates a stationary acoustic pressure field in the chamber: two nodes are typically located at 25 *μ*m and 75 *μ*m altitude in the 100 *μ*m high fluidic channel (Fig. 1) [17]. Starting from a value pre-calibrated by the manufacturer for each AFS chip, the driving frequency was adjusted manually in order to reach the so-called resonance frequency (RF) for which the vertical pressure gradient is maximized at the vicinity of the chamber floor. In case of force drop due to RF drift, for example caused by minute temperature change, the frequency was slighly readjusted by the user in order to increase the force before starting a new measurement cycle. The driving tension amplitude, called a ‘power’ by the manufacturer, is expressed in % units, and is said to scale linearly with the force applied to a bead at a given altitude.

The temperature inside the chamber is regulated to 25°C by a proportional–integral-derivative (PID) controller, a sensor being placed in the middle of the flow cell and two heating elements warming up the whole flow cell.

### Measurement protocol and data acquisition

The setup is controlled through the Labview software provided by the manufacturer (Lumicks, v1.4.0), which we have modified for our needs. More precisely, we have added an option to generate periodic acoustic force patterns corresponding to tunable steps of power *P* and duration *T*. For example, in a typical unbinding experiment, a 0% power period of typical duration *T*^*A*^ ≥5 min is used to determine the anchor point of each bead. Next, a cycle alternating a very low power (*P*^*L*^ *<* 0.01%) period and test power period (power *P*) is repeated typically 100 times. The corresponding durations *T*^*L*^ and *T* need to be adapted to the nature of interacting partners, and can vary from a fraction of a minute up to 15 min. As a rule of thumb, *T* can be set as at least 2 or 3 times the inverse of the expected off-rate.

The AFS software also allows one to track each bead in 3D on the fly thanks to real-time image. The x,y position corresponds to the center of the bead diffraction pattern, while its relative *z*-position is determined using a predefined look-up-table (LUT) ranging from 0 nm to 10 000 nm in 100 nm steps [19]. The individual bead traces are recorded in a tdms file (Labview format), together with the power amplitude, frequency, and temperature.

### Data analysis

The data analysis is performed in two phases: a) during steps 1-6 described below, one single tdms file is analyzed. It contains cyclic measurement data at a single test power of multiple tethered beads from one field of view. b) during steps 7-8, the outcome of the first phase for various measurements at different test powers is merged for each individual beads. Global fitting procedures are performed and synthesis plots are produced.

#### 1. Initialization

The tdms file contains the bead traces (x,y,z coordinates vs time) acquired from one field of view during one cyclic measurement (in this case the procedure is used in ‘cycle’ mode). One or several test traces are selected for analysis as well as several reference traces (if available). Test traces are typically obtained from beads tethered via a single functional J-DNA. Reference traces are typically obtained from beads directly bound to the surface, exhibiting minimal fluctuations and allowing to correct for drift of the chamber position over time. The time interval for the analysis of all traces is also set.

The following steps 2-6 are applied for each test trace.

#### 2. Baseline correction

This operation compensates for apparent drift of the sample due to lateral and vertical motion, caused for example by the dilatation of the microscope elements under temperature variation of the environment. The correction is performed in 3 directions by calculating an average reference trace mimicking the motion of the whole sample and withdrawing it from the test trace. If several reference traces are available, the average reference trace is calculated as the mean for each coordinate, followed by a rolling average over typically 100 s. If no surface-bound bead is available for reference, a self-reference can be performed on a J-DNA tethered bead. The self-reference trace is built taking as a basis the coordinates from the test bead as follows: the *z* position of the test bead is selected at low power, while the *x, y* positions of the test bead are selected at high power. The rest of the trace is obtained by interpolating by a constant value in order to fill in the gaps at high power for *z* and low power for *x, y*. The complete self-reference trace is obtained after a rolling average on 100000 time points, in order to completely smooth out the effect of interpolation.

#### 3. Anchor point determination

One here relies on the data acquired during the dedicated time interval *T*^*A*^ or on all time points for which *P* = *P*^*L*^. The anchor point coordinates (*x*_A_, *y*_A_, *z*_A_) are obtained, in the horizontal plane, as the time average of *x, y* and, along the vertical axis, as the minimal value of *z* after a rolling average on 400 points (Fig. S2). In case the bead is not exploring the whole available space, for example due to transient binding to the surface, the center of the envelope of the *x, y*-positions is used instead of in-plane barycenter (Fig. S2B). For this, a kernel density estimation of the *x, y*-coordinates distribution is performed, defining typically 20 contour lines, and the center of the external contour is taken for setting (*x*_A_, *y*_A_).

#### 4. Tether length and pulling angle determination

The length *L* of the scaffold corresponds to the distance between its point of attachment to the bead and the point of anchoring A to the surface (Fig. S2). Noting *x, y, z* the coordinates of the point of lowest altitude at the surface of the bead, one has:

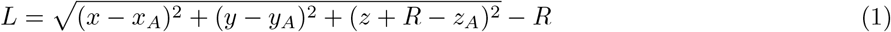

where *R* is the radius of the bead. The pulling direction was determined by calculating the polar angle *θ* (Fig. S2):

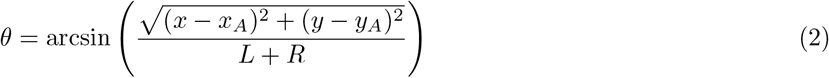

#### 5. Opening events detection

This operation is performed on the *z − z*_*A*_ trace, separately on each time interval during which the acoustic power is applied. A 1D gaussian filter of size = 36 points is first applied and the signal is normalized between 0 and 1. The crossings of a 0.5 threshold value are recorded and the step size is determined, following the original procedure found in github.com/thomasbkahn/step-detect. If less than 5 steps are recorded for a given high power interval, the step largest amplitude is taken for the opening transition. If more than 5 steps are recorded, the interval is considered as featuring no opening events. In this case, the average extension is compared with the middle altitude *Z*_m_ representing the middle between open and closed state, typically 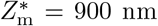. If 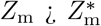, one considers that the J-DNA did not close back at the previous low power interval. If 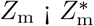, one considers that the J-DNA did not open during the test power interval. The lifetime of the close state is measured between the onset of high power application plus a delay and the opening time. The delay accounts for the time to reach the effective force on the bond, and empirically set to 1-2 s.

#### 6. Power spectrum calculation

The single sided power spectrum density (PSD) of the signal is calculated separately on *x − x*_*A*_ and *y − y*_*A*_ thanks to the signal.periodogram routine from Python Scipy package. In a default mode that is used for the ‘force calibration on the fly’, we take into account all the points tagged as belonging to the open state. Alternatively, in ‘calibration mode’, a specific time interval can be selected, for which steps 1-4 are performed as above and step 5 is skipped. This mode can also be used to calculate the PSD using data obtained at a high frame rate and specifically dedicated for calibration of large forces. For graphical representation, PSD are binned on an equally spaced log frequency scale.

#### 7. PSD fitting and force calibration

For a 55 Hz acquisition frame-rate, we typically work with frequency *f* ranging from 0.1 to 15 Hz. For a given bead, the PSD along *x* at power *P*^*i*^ is compared with [62]:

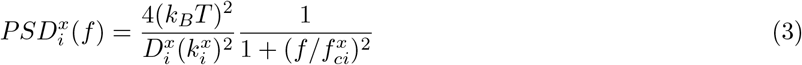

where 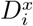 is the bead diffusion coefficient parallel to the surface, 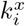 the stiffness of the pendulum, and 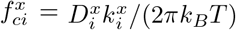 the corner frequency, delineating the elastic and viscous parts of the spectrum obtained at power i. *k*_*B*_*T* is the thermal energy at temperature *T*.

Writing *z*_*i*_ the bead height from the chamber surface at power *P*^*i*^ and *R*^*x*^ the bead radius, the diffusion coefficient can be approximated by [63]:

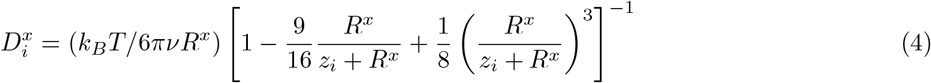

where *ν* is the viscosity of water. A global fit using Eq. 3 and Eq. 4 is then applied to all 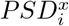 simultaneously, taking the 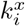 and *R*^*x*^ as fitting parameters. Alternatively, individual PSD fit can also be performed applying Eq. 3 and taking the 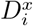 and 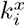 as fitting parameters. This is possible when the condition 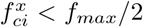 is fulfilled, i.e. for low power or high acquisition frequency. The force determined along *x* for a given bead at power *P*^*i*^ is given by:

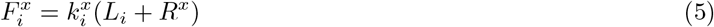

where *L*_*i*_ is the scaffold extension in its open state. The average vertical force and bead radius taking into account the analysis one can perform in both directions are simply calculated as : 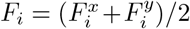 and *R* = (*R*^*x*^ + *R*^*y*^)/2.

The force applied along the J-DNA when pulling at angle *θ* is given by:

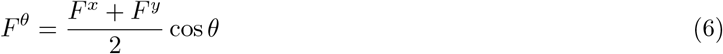

#### 8. Survival curves and fitting

For a given bead and a given power, all the dwell-times before opening, Δ*t*, are pooled. It also includes the no-open events to which we attribute *T* as dwell time, i.e. the duration of application of the acoustic power. All those values are next sorted by increasing order so as to build a survival fraction S corresponding the fraction of J-DNA closed until time t as a function of t. The survival curve is fitted with a monoexponential function which appears on a semilog graph as a straight line with a slope equal to the bond off-rate. The uncertainty on the off-rate is calculated as the difference of slope of the survival curves calculated either on the first 50 unbinding events or the last 50 unbinding events.

## Results and Discussion

### Principle of the coupling of DNA scaffolds and AFS

After the beads have been injected in the AFS chamber and allowed to settle for a few minutes, the application of a limited acoustic power (1%) permits to collect the non-tethered ones at the first pressure node along *z*, far out of focus, and to flow them away. One can thus select the beads that are tethered to the surface via J-DNA molecules (Fig. 1A) and track them using the acquisition software (Fig. 1B). Additional sorting can be performed by eliminating the beads for which the recorded *x,y*-trace is strongly anisotropic, which is a typical feature of beads tethered via multiple J-DNA. If available in the field of view, one or several beads showing no Brownian motion can be selected as reference beads to enable baseline drift correction. Recording of the look-up tables (LUT) that contains, for each bead, its diffraction pattern as a function of the defocusing is then performed by applying an acoustic power of 1 % and moving the microscope stage in the *z* direction (Fig 1B) [19, 17].

In a typical measurement the piezo is driven by a rectangular wave, the acoustic power being periodically switched between two values: one nearly equal to zero and one able to put the scaffold under tension (Fig. 1C). Starting at low force with the proteins dissociated, the bead is at low altitude and it exhibits strong fluctuations (Fig. 1D, left part of height profile in grey): the molecular partners can easily access each others but one can not distinguish when binding occurs. Upon application of a 2% test acoustic power, the height quickly increases to a plateau value and exhibits reduced fluctuations, corresponding to the closed state of the scaffold with protein complex formed (Fig. 1D, orange trace). A sharp transition to a higher plateau is next observed, corresponding to the transition to the open state where the complex is dissociated (Fig. 1D, red trace). The difference between the two plateaus corresponds to approximately 200 nm, as expected from the length of the J-DNA leash (Table S2). The acoustic power is finally reduced, releasing the tension on the leash and permitting rebinding (Fig. 1D, right part of trace in grey).

Once the *x,y,z* traces corrected for drift and the J-DNA anchor points at the chamber surface determined, one proceeds to the analysis of the cyclic experiment *per se*. An example of *z*-trace obtained during 100 power cycles is shown on Fig. 2A with a zoom provided in Fig. 2B. An automatic step detection permits to : 1) measure the scaffold length extension upon jump (see Fig. 2B); 2) distinguish the open and closed states on the trace at test power, as represented by the orange and red colors in Fig. 2A,B. 3) measure the duration Δ*t* in closed state (bound complex) under application of high power (see Fig. 2B). The corresponding survival curve obtained from Δ*t* is shown in Fig. 2C. The density distribution of Δ*z* jumps is represented in 2D. The values are peaking at around 200 nm, as expected from Table S2. Similar plots acquired for other values of the acoustic power evidence a decrease of the bond lifetime and an increase of jump amplitude with the applied force.

**Figure 2:**
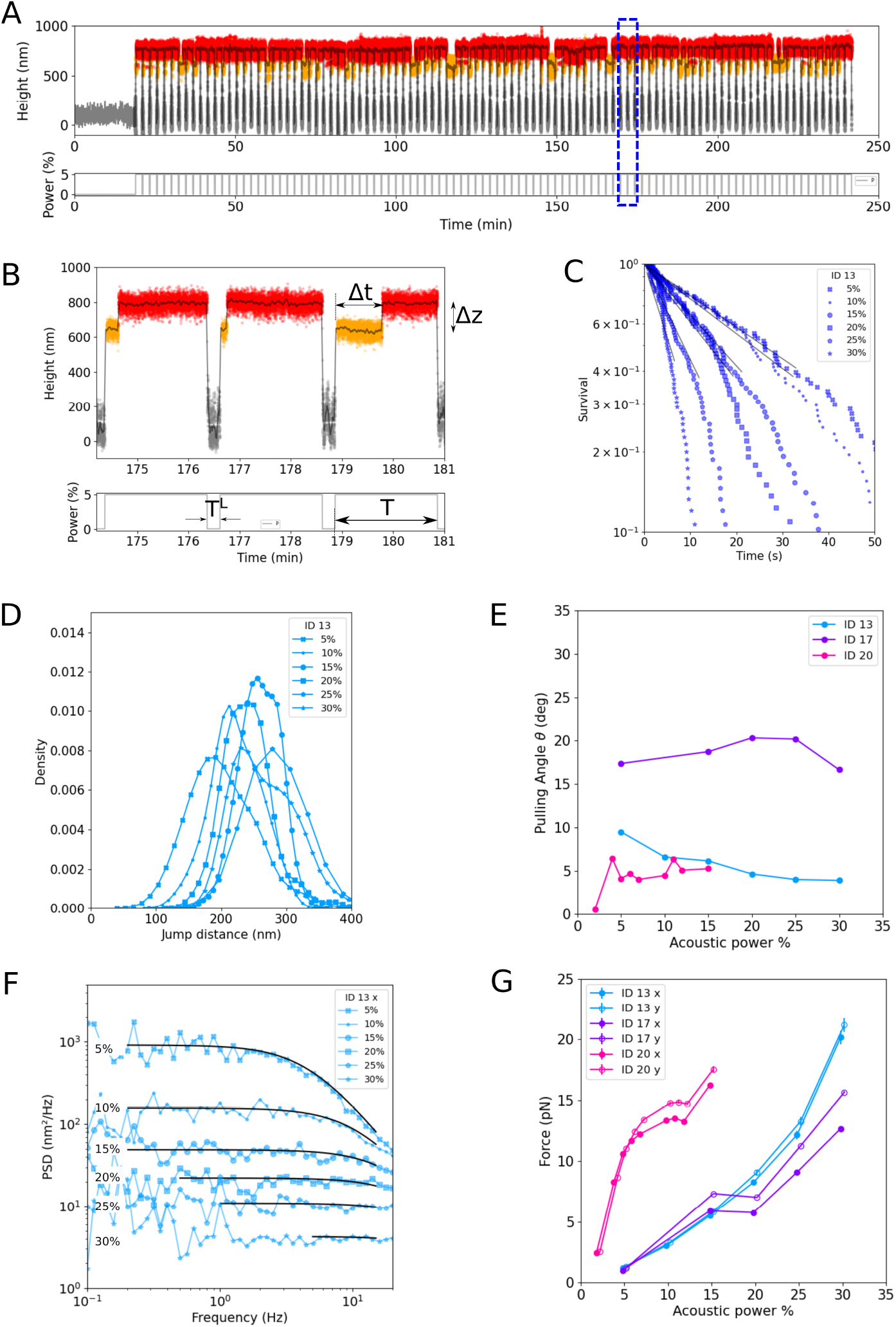
Tracking of a single bead under force cycles, bond lifetime determination and force calibration. (A) Vertical trace *z* obtained during 100 alternate low/high acoustic power cycles. (B) Zoom of trace in A, indicated by the dashed blue rectangle. Based on power value and jump detection at high power, the trace is divided in 3 subsets: low power (grey), closed state under force (orange), open state under force (red). The lifetime Δt of the bond corresponds to the duration of the orange step. Δz represents the height jump between closed and open state. (C) Survival curve calculated from the lifetime Δt distribution for one representative bead at different acoustic powers in %. Superimposed mono-exponential fits : *S*(*t*) = exp(Δ*t*.*k*_off_). (D) Density kernel of jump distance Δz for the same bead as in (C) at different acoustic powers. (E) Pulling angle vs acoustic power for 3 representative beads. (F) Power spectrum density for the same bead as in D, at various applied powers i. Curves fitted to determine bead radius *R* and stiffness *k*_*i*_ are shown as black lines. (G) Force vs acoustic power for 3 representative beads, the same as in E. Plain symbols from x direction. Empty symbols from y direction.

### Pulling angle measurement

The pulling angle *θ* is the angle formed between the *z*-axis and the line passing by the anchor point and the bead center (see Fig. S2 for a geometrical depiction with the open J-DNA and Eq. 2 for the way this parameter is obtained from the measurements). As can be clearly seen in fig. 2E for three representative beads *θ* does not usually depend on the applied power. We also observed that in a same field of view its average value varies between 0 and 30° (data not shown). However, more strikingly, the pulling angle for a same bead may sometimes increase or decrease continuously by nearly 30° over the explored power range. This illustrates the existence of lateral pressure nodes which orient the pulling axis and which position can change with power [40, 41].

### Force calibration and scaffold length measurement

The acoustic force applied to each bead is determined by calculating the power spectrum density (PSD) and relating it to the stiffness of an inverted pendulum having the length of the scaffold [19, 39], following Eq. 5 (see Materials and Methods). We first examined the PSD and the way to fit them.

#### PSD computation

The PSD are computed from the bead lateral fluctuations for each test power. A typical example obtained in the *x* direction is represented on Fig. 2F. Traditionally, these data are obtained separately by applying a constant power for a given time. We propose to extract these data directly from the applied force cycles (see Materials and Methods). This calibration “on the fly” avoids adding an additional calibration step. To show that this method provides the same results than a “single step calibration”, we performed both at the same test powers on the same scaffolds. Fig. S3 shows that the PSD are almost identical. The ‘on-the-fly’ method provides a less noisy signal at low frequency since the signal has been accumulated over a longer time in total.

#### PSD fitting

We then propose to perform a global fit, using Eq. 3 and 4, considering all the tested powers for each bead, and determining a single bead radius and a value of stiffness *k* for each tested power. In order to validate this global fit method, we have alternatively fitted individual PSDs at different power *P*^*i*^ for diffusion *D*_*i*_ and stiffness *k*_*i*_. The obtained *D*_*i*_ values obtained are independent of the applied power and match those deduced from the global fit (see one example on Fig. S4). This validates the global fit method, which allows to overcome some individual fit difficulties, when the corner frequency cannot be pinpointed on the PSD obtained with the low frequency acquisition utilized in the cyclic measurement. By combining “on the fly” calibration and global fit, we were able to compute and fit the PSD for all beads at all tested powers. The results for 21 beads tethered by J-DNA functionalized with the FKBP12 and FRB complex are shown on Fig. S5 and Fig. S6 for *x* and *y* directions respectively. The fits are superimposed to the data showing a good agreement for all powers. In addition, PSDs from 9 beads functionalized with the Nef and Nef19 are shown on Fig. S7, also showing a good fit quality.

#### Fitted bead radius

The distribution of *R*^*x*^ or *R*^*y*^ for both molecular complexes fairly agrees with the nominal values provided by the manufacturer (Fig. S8). However, we systematically observed *R*^*x*^ *< R*^*y*^, which we attributed to a poorer data and fit quality in the *y*-direction, possibly related to the fact that the rectangular AFS chamber (smaller in *y* direction and therefore more sensitive to lateral reflection phenomena in this direction).

#### Fitted stiffness

Fig. S9 shows the fitted stiffnesses for different FKBP12-FRB scaffolds and powers. The stiffnesses measured in *x* or *y* directions exhibit very similar values. A non-linear dependence with applied power is also noted, the curves being either concave or convex. This could be due to minute shift of the resonance frequency away or towards the driving frequency.

#### Length of J-DNA

The length of the J-DNA is measured between the anchor point at the chamber surface and the attachment point onto the bead, as illustrated by Fig. S2A (see Eq. 1 for its computation). The variation with acoustic power of the length of open J-DNA is shown for FKBP12-FRB scaffolds in Fig. S10. The maximum extensions often exceed the expected scaffold length listed in Table S2. One can here invoke some variation due to the random integration of digoxygenin and biotin at the extremities of the shanks. It also possibly points to a limited precision in the determination of the anchor point position. Some improvement could here be made by application of a lateral flow in the chamber in two opposite directions and consider the middle of the two extreme bead positions. However, it is not possible to supercoil the tether to bring the bead on the anchor point as done with dsDNA and MT.

#### Force calculation

The force is then calculated following Eq. 5. Similarly to the stiffness, it depends non-linearly on the acoustic power (see Figs. 2G, S11). More specifically, the force typically varies from 0 to 20 pN as the acoustic power is turned up. However, both shape of the response and maximum value one can achieve seems to vary significantly between different positions in the chamber, once more illustrating the existence of acoustic nodes in *x* and *y* [41]. The variation with force of the length of open J-DNA is shown for FKBP12-FRB scaffolds in Fig. S12. While, as expected, the length monotonously increases with applied force in most cases, some curves exhibit non-monotonic behaviour. This could stem from a limited precision in extended state determination due to errors in jump determination. Also, we did not succeed in satisfactorily adjusting the data to either the freely jointed chain model or to the worm-like chain one. This is possibly because of the lack of data collected at very low force. In fact, to obtain a more accurate determination of the scaffold mechanical properties, dedicated investigations in the low force domain would be required, which are out of the scope of the present paper.

### Force dependence of the off-rate

The use of J-DNA allows repetitive measurements on individual bonds, cycling between the association of the proteins when the acoustic power is low and their dissociation when it is high (Fig. 1C and 2A).

The fraction of non-closed scaffolds observed when increasing *P* remains usually less than 0.2, indicating that the duration *T*^*L*^ the molecules spend at low force is sufficient for an efficient rebinding. It was also noticed that, for beads of 1.58 *μ*m in diameter, the cloud of points at low force exhibits a symmetric and homogeneous distribution around the anchor point (Fig. S2B), which we relate to a correct scaffold recoiling and thus more opportunities for the reactive partners to encounter.

At the test acoustic power, one measures the dwell time Δ*t*, which corresponds to the time elapsed until an extension jump Δ*z* is detected (Fig. 2B). The dwell times collected over the force cycles are pooled in a survival curve, which is established for each J-DNA and each applied force. These survival curves are individually fitted by a single mono-exponential function, yielding a straight line which slope is the off-rate when represented in semi-log scale (Fig. 2C and S13). Such off-rates *k*_off_ obtained for the FKBP12-rapamycin-FRB complex are next plotted as a function of applied force *F* for each scaffold and fitted separately with the Bell model:

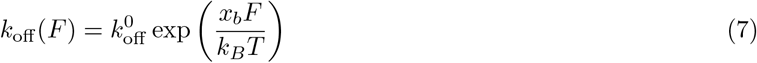

where 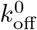 is the off-rate in the absence of applied force and *x*_*b*_ is the distance from the potentiel well to the barrier on the interaction energy landscape (Fig. S14). Since the existence of lateral forces have been evidenced, to ensure that the influence of the pulling angle remains negligible, we computed the force projected along the pulling angle (Eq. 6). The Bell parameters obtained considering these forces differ only marginally from the ones determined without projection of the force (see Fig. S14 for a comparison).

A superposition of all Bell plots illustrates the limited dispersion of the data (Fig. 3A). As also evidenced on Fig. 3B, individual Bell analyses performed on each of the 21 J-DNA functionalized with FKBP12 and FRB provide 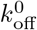 and *x*_*b*_ values which dispersion remains very limited (¡60%) and in the same range to what is observed between publications (see also Fig. S14 as well as Tables S3 and S4) [22, 30, 45, 46]. Alternatively, all *k*_off_ values are pooled by force bins (Fig. 3A, black disks) and the data were fitted using a single Bell curve: it gave 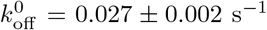 and *x*_*b*_ = 0.42 *±* 0.04 nm. As for the individual fits, these results are in good agreement with the ones retrieved with the same J-DNA mounted on magnetic tweezers: 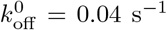 and *x*_*b*_ = 0.42 nm [30] (Table S4). The off-rates extrapolated at zero force are also consistent with the determination achieved by SPR, 20.5× 10^−3^ s^−1^ [46] and 16.7× 10^−3^ s^−1^ [45] (Fig. 3A, B and Table S3), as well as by SMFS, 26 and 32 × 10^−3^ s^−1^ [22].

**Figure 3:**
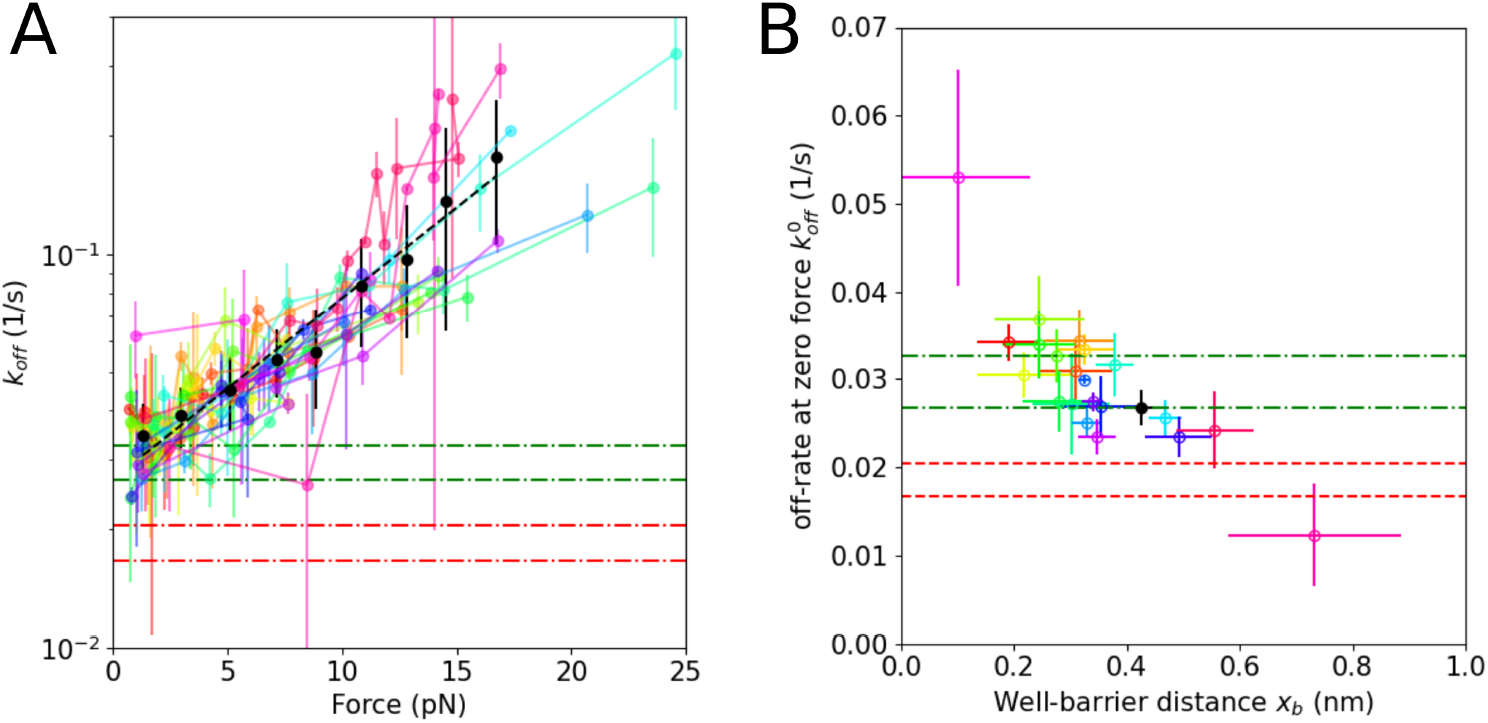
Bell plots for the FKBP12-rapamycin-FRB complex and fit. (A) The off-rate at different forces of individual molecules is shown in different colors. The error bars on invidual off-rate values are calculated as indicated in the Materials and Methods. The average values on force bins of equal size are shown as black disks. The vertical error bars on the black points are calculated as the standard deviation of off-rate values in each force bin. The dashed black line is the fit with Bell equation 7. (B) Off-rate in absence of applied force 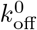 as a function of *x*_*b*_, the distance from the potential well to the barrier on the energy landscape of the interaction, as obtained from individual Bell fits shown in Fig. S14. The error bars are the fitting errors when applying Bell model. The dashed lines in A and B represent the values of off-rate measured by Surface Plasmon Resonance [45, 46] and SMFS [22, 30].

The same measurements and analyses were performed for 9 J-DNA scaffolds functionalized with Nef and Nef19. This complex being much more long-lived than FKBP12-rapamycin-FRB, we aimed at applying higher forces to achieve unbinding in reasonable times. We used larger beads (3*μ*m diameter), in order to increase the acoustic force at equal power, without generating too much heat or shifting the resonance too strongly. The force vs acoustic power data are represented on Fig S15: a sharp force increase can be observed when changing the diameter of the beads from 1.58 to 3 *μ*m. However, as displayed on Fig. S16 survival curves, despite that force increase, the bond lifetime often exceed the measurement time *T*. Thus, survival curves contain a limited number of rupture events and are censored at a maximal duration between 600 and 1200 s. While, due to this limitation, independent curve fit of off-rate vs force are not accurate (data not shown), one can perform a fit on pooled data, as shown on Fig. 4. The Bell parameters are as follows: 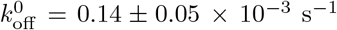 and *x*_*b*_ = 0.75 *±* 0.08 nm. As for FKBP12-rapamycin-FRB, there is a satisfactory match with the 0.18× 10^−^ s^−1^ off-rate determined by SPR in the absence of force [55]. Notice that *x*_*b*_ is almost doubled compared to the case of FKBP12-rapamycin-FRB, while 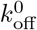 is reduced by a factor 10. Thus the Nef-Nef19 bond is much more stable at low force, but more sensitive to force.

**Figure 4:**
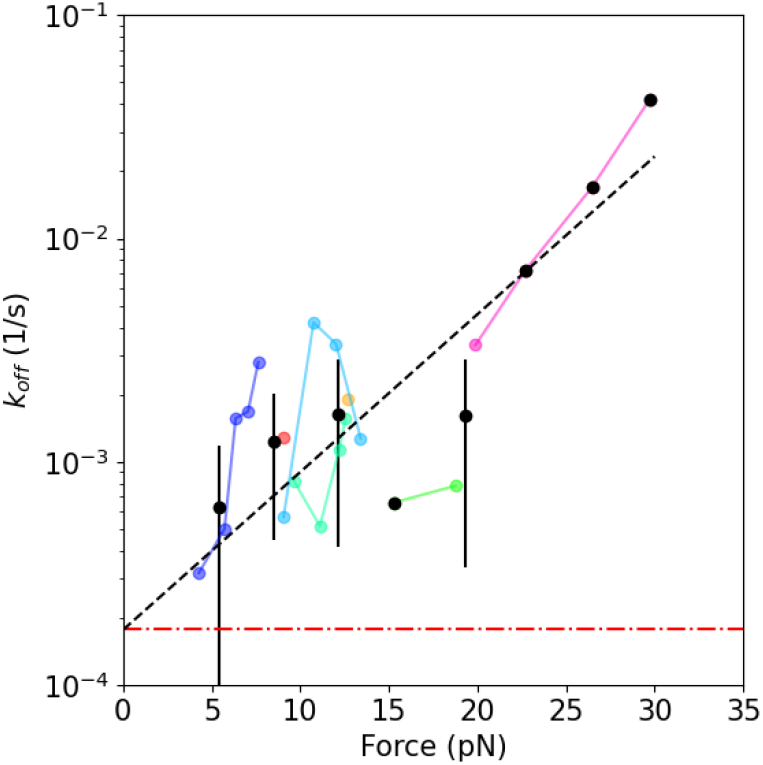
Bell plot for the Nef-Nef19 complex. The black points correspond to the *k*_off_ averaged by equal force bins. The dashed black line is the fit with Bell equation 7. The vertical error bars on the black points are calculated as the standard deviation of off-rate values in each force bin. The horizontal dashed line represents the value of the off-rate measured by SPR [55].

## Discussion

AFS is a parallel SMFS technique that gives potential access to a larger force range than its competitors. However, a practical constraint lays in the current commercial closed format of the chambers, for which the development of regeneration procedures seems essential in order to guarantee the reproducibility of bead tethering. Besides, as for other parallel methods like laminar flow chamber and MT, a trade-off for the choice of bead size should be found to optimize force range while limiting non-specific adhesion. A second drawback resides in the strong heterogeneity and the limited stability of the acoustic field. Thus, one can not expect to map the force field in a preliminary experiment and rely on the collected informations to perform subsequent investigations. In fact, force calibration has to be realized for each scrutinized protein complex. J-DNA offer this opportunity because they are both scaffolds on which the molecular partners can be engrafted and tethers that can be used for force determination by the inverted pendulum method. Force calibration is achieved by collecting the calibration data “on the fly” during the measurements and fitting them globally for each J-DNA, which allows maximum benefit to be derived from data acquired under standard experimental conditions at regular acquisition frequency. Interestingly, such *in situ* calibration is hard to enforce with MT for instance, the orientation imposed to the bead by the magnets impeding accurate length measurement for DNAs only several kilobase pair-long.

Hence, the present work constructively combines AFS and J-DNA to study the response to force of single biomolecular bonds with individual calibration of the applied force. Utilizing a scaffold enables to observe enough dissociation events on the same two protein partners so that one can probe bond aging, a phenomenon we have not observed so far, the off-rate calculated either during the first 50 cycles of a measurement or during the last 50 ones being essentially identical (Fig. S17). It also enables to extract the individual Bell’s parameters for individual protein complexes. As a consequence, one can now compare the different complexes and challenge the concept of bond heterogeneity, as formalized in [64]. Interestingly, our setup removes some of the possible environmental causes of heterogeneity, which are listed in [64], including variable attachment of proteins to surface and non-specific interactions. We observe for the FKBP12-rapamycin-FRB complex a limited dispersion of the Bell parameters, as compared to variations between publications from the literature, as well as a correlation between 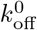 and *x*_*b*_, which is discussed below. Overall, these observations suggest a limited heterogeneity of the FKBP12-rapamycin-FRB complex. Homogeneity has indeed been recently measured for some other protein unfolding [65]. We finally remark that the absence of bond heterogeneity may not question the concept of roughness of the energy landscape [66, 67, 68].

Examination of Fig. 3B shows a negative correlation between 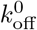 and *x*_*b*_ for the FKBP12-rapamycin-FRB complex. To assess where this trend could originate from, we first hypothesized a role for the visco-elastic properties of the experimental system [69]. However, we observe no correlation between the *x*_*b*_ parameter and mechanical parameters of the scaffold, such as the J-DNA length and elasticity, or the bead size (data not shown). Additionally, we computed the effect of a parabolic potential imposed by the spring-like scaffold, as proposed in [70], and found that the quadratic term induces a negligible increase of *k*^0^ for the highest *x*_*b*_ measured. Another possibility to explain Fig. 3B correlation is the involvement of the curvature of the energy landscape, which predicts a dependence of 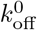 as 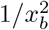, as proposed for example in [71]; however this scaling with *x*_*b*_ is too strong compared to our data (not shown). Finally, we propose that the error in fitting the slope in the *k*_off_ vs F plots leads to a correlated error in the intercept of the *y*-axis. Further refinements of the present setup, in particular for anchor point determination, stability of the setup, precision of jump detection as well as the precision of force calibration would certainly help to address this issue.

As a conclusion, our technique has a sufficient accuracy to characterize the force-induced rupture of individual FKBP12-Rapamycin-FRB and antigen-nanobodies complexes. We also illustrate how the method can allow the comparison of individual bonds of same nature and bond behaviour under repeated actuation. Therefore, it has the potential to both answer fundamental questions and open the way to systematic chemomechanical characterization of biomedically relevant biomolecular interactions.

## Author Contributions

CG and LL designed the research. JYW and CV carried out experiments. AA, PC, MF, DK, TS, CG contributed reagents. LV and FR contributed to the acquisition software and to the experimental setup. JYW, CV, FR, and LL analyzed data. JYW, CV, CG, and LL wrote the manuscript. All authors critically revised the manuscript.

## Acknowledgments

We thank Lumicks (in particular Douwe Kamsma and Eugen Ostrofet), as well as Pierre-Henri Puech for fruitful discussions. The AFS setup was acquired thanks to the grant Projet exploratoire région PACA 2017 – AcouLeuco obtained by FR. We thank AMIDEX Emergence Innovation project ForSelecAntibody, Plan Cancer PhysCancer program (project ComPhysAb), European Research Council (ERC, grant agreement No. 772257), Human Frontier Science Program (HFSP, grant No. RGP0056/2018), PSL valorisation (Grant J-DNA), Marie Curie Sklodowska action (MSCA-IF, grant agreement 895819), and Labex Memolife. The Molecular Motors and Machines team at IBENS is an “Equipe labellisée” by the Ligue Nationale Contre le Cancer. We also acknowledge INSERM, CNRS and AMU for regular support.

## Supplementary Material

### Supplementary Tables

**Table S1:** Geometrical parameters characterizing the various anatomical segments making up the two J-DNAs used in the present study (see Fig. S1A for a pictorial description). The asymmetrical scaffold is the one described in Kostrz et al. [1]; the symmetrical one has been obtained following the same synthesis protocol except that the sequences of the TS_1_ and TS_2_ oligonucleotides were respectively changed for ATATGAG<u>GCTGAGG</u>GCAGCCACTGGTAACAGGATTAGCAGAGCGAGGFATGTAGGCGGTGCTACAGAG and TGTAAGA<u>GCTGAGG</u>TCGCAATGGAGTGTCATTCATCAAGGACGCCGCFATCGCAAATGGTGCTATCC (5’ to 3’ direction, F = azido-dT, underlined bases correspond to the Nb.BbvCI nicking site). The extension jumps associated with the detection of a protein-protein interaction rupture were computed by subtracting the length of the scaffold in the closed state to the one in the open state.

| J-DNA | Leash | Shank 1 | Tip 1 | Shank 2 | Tip 2 | Shanks + Tips<br>(closed scaffold) | Shanks + Leash<br>(open scaffold) | Extension<br>jump |
| --- | --- | --- | --- | --- | --- | --- | --- | --- |
| <u>Contour length (bp)</u> |  |  |  |  |  |  |  |  |
| Asymmetrical | 689 | 629 | 48 | 1 348 | 42 | 2 067 | 2 666 | 599 |
| Symmetrical | 689 | 1 459 | 48 | 1 422 | 48 | 2 977 | 3 570 | 593 |
| <u>Contour length (nm)</u><sup>a</sup> |  |  |  |  |  |  |  |  |
| Asymmetrical | 233 | 213 | 16 | 445 | 14 | 699 | 901 | 202 |
| Symmetrical | 233 | 493 | 16 | 481 | 16 | 1 006 | 1 207 | 200 |

<sup>a</sup> Calculated assuming a rise of 3.38 Å per base pair [2].

**Table S2:** Geometrical parameters characterizing the different transitions one can observe with the symmetrical J-DNA scaffold. There is only one maximal extension conformation, which identifies with the open state, but there are five possible lower extension conformations, which correspond to the two proteins specifically interacting together in the closed state or to one of the proteins interacting with one of the surfaces (see Fig. S1C for a pictorial description). The contour lengths of the various anatomical segments were selected so as to associate a unique extension jump value with the detection of a protein-protein interaction; hence, all the other step variations, which are larger, report on non-specific phenomena.

| Interaction | Segments separating<br>the chamber surface from the bead one | Contour length (bp) | Contour length difference with<br>the open state (bp) |
| --- | --- | --- | --- |
| <u>High extension conformation</u> |  |  |  |
| None | Shank 1 + Leash + Shank 2 | 3 570 | 0 |
| <u>Low extension conformations</u> |  |  |  |
| Protein 1 – Protein 2 | Shank 1 + Tip 1 + Tip 2 + Shank 2 | 2 977 | 593 |
| Protein 1 – Surface 1 | Tip 1 + Leash + Shank 2 | 2 159 | 1 411 |
| Protein 2 – Surface 2 | Tip 2 + Leash + Shank 1 | 2 196 | 1 374 |
| Protein 1 – Surface 2 | Tip 1 + Shank 1 | 1 507 | 2 063 |
| Protein 2 – Surface 1 | Tip 2 + Shank 2 | 1 470 | 2 100 |

**Table S3:**
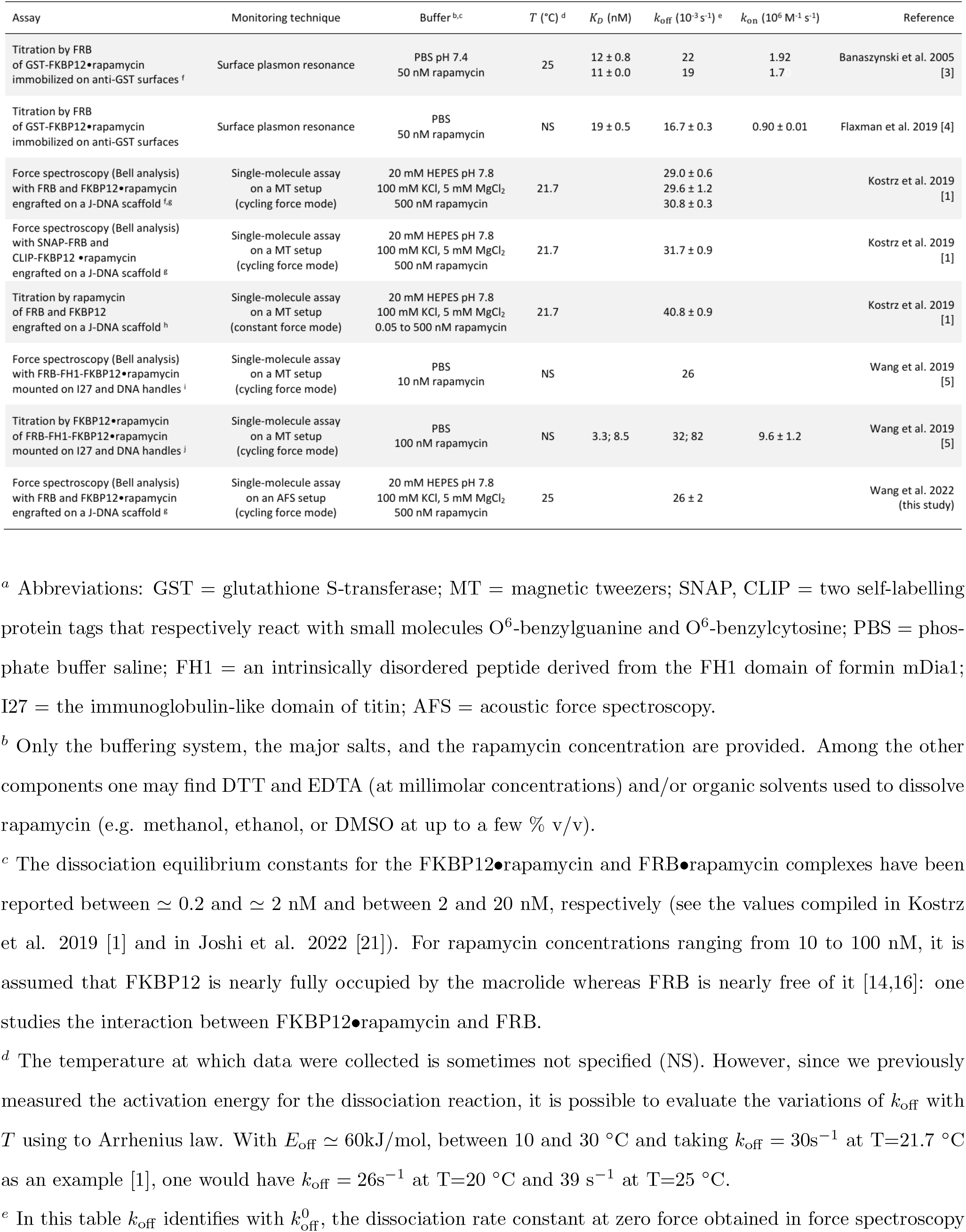
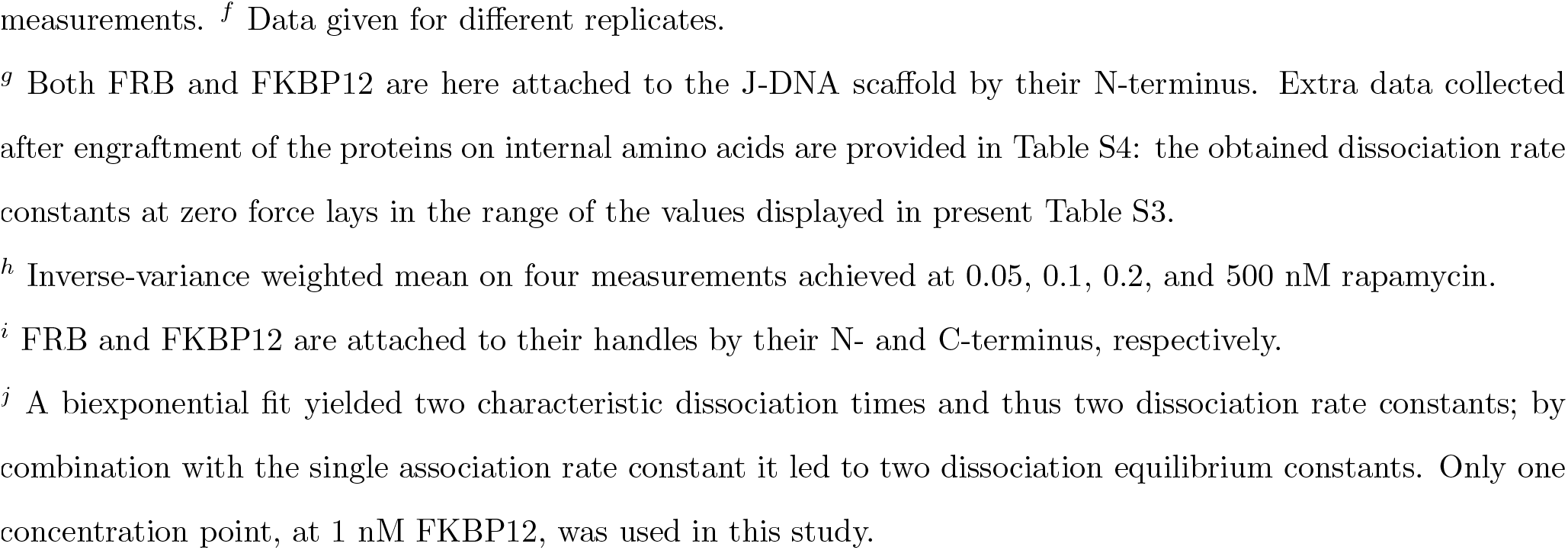
Published kinetic parameters for the FKBP12•rapamycin•FRB *⇌* FKBP12•rapamycin + FRB reaction. *K*_*D*_ corresponds to the dissociation equilibrium constant, *k*_off_ to the dissociation rate constant, and *k*_on_ to the association rate constant. ^*a*^

| Assay | Monitoring technique | Buffer <sup>b,c</sup> | $T$ (°C) <sup>d</sup> | $K_D$ (nM) | $k_{\text{off}}$ ($10^{-3} \text{ s}^{-1}$) <sup>e</sup> | $k_{\text{on}}$ ($10^6 \text{ M}^{-1} \text{ s}^{-1}$) | Reference |
| --- | --- | --- | --- | --- | --- | --- | --- |
| Titration by FRB of GST-FKBP12•rapamycin immobilized on anti-GST surfaces <sup>f</sup> | Surface plasmon resonance | PBS pH 7.4<br>50 nM rapamycin | 25 | $12 \pm 0.8$<br>$11 \pm 0.0$ | 22<br>19 | 1.92<br>1.7 | Banaszynski et al. 2005 [3] |
| Titration by FRB of GST-FKBP12•rapamycin immobilized on anti-GST surfaces | Surface plasmon resonance | PBS<br>50 nM rapamycin | NS | $19 \pm 0.5$ | $16.7 \pm 0.3$ | $0.90 \pm 0.01$ | Flaxman et al. 2019 [4] |
| Force spectroscopy (Bell analysis) with FRB and FKBP12•rapamycin engrafted on a J-DNA scaffold <sup>g</sup> | Single-molecule assay on a MT setup (cycling force mode) | 20 mM HEPES pH 7.8<br>100 mM KCl, 5 mM MgCl<sub>2</sub><br>500 nM rapamycin | 21.7 | | $29.0 \pm 0.6$<br>$29.6 \pm 1.2$<br>$30.8 \pm 0.3$ | | Kostrz et al. 2019 [1] |
| Force spectroscopy (Bell analysis) with SNAP-FRB and CLIP-FKBP12•rapamycin engrafted on a J-DNA scaffold <sup>g</sup> | Single-molecule assay on a MT setup (cycling force mode) | 20 mM HEPES pH 7.8<br>100 mM KCl, 5 mM MgCl<sub>2</sub><br>500 nM rapamycin | 21.7 | | $31.7 \pm 0.9$ | | Kostrz et al. 2019 [1] |
| Titration by rapamycin of FRB and FKBP12 engrafted on a J-DNA scaffold <sup>h</sup> | Single-molecule assay on a MT setup (constant force mode) | 20 mM HEPES pH 7.8<br>100 mM KCl, 5 mM MgCl<sub>2</sub><br>0.05 to 500 nM rapamycin | 21.7 | | $40.8 \pm 0.9$ | | Kostrz et al. 2019 [1] |
| Force spectroscopy (Bell analysis) with FRB-FH1-FKBP12•rapamycin mounted on I27 and DNA handles <sup>i</sup> | Single-molecule assay on a MT setup (cycling force mode) | PBS<br>10 nM rapamycin | NS |  | 26 |  | Wang et al. 2019 [5] |
| Titration by FKBP12•rapamycin of FRB-FH1-FKBP12•rapamycin mounted on I27 and DNA handles <sup>j</sup> | Single-molecule assay on a MT setup (cycling force mode) | PBS<br>100 nM rapamycin | NS | 3.3; 8.5 | 32; 82 | $9.6 \pm 1.2$ | Wang et al. 2019 [5] |
| Force spectroscopy (Bell analysis) with FRB and FKBP12•rapamycin engrafted on a J-DNA scaffold <sup>g</sup> | Single-molecule assay on an AFS setup (cycling force mode) | 20 mM HEPES pH 7.8<br>100 mM KCl, 5 mM MgCl<sub>2</sub><br>500 nM rapamycin | 25 | | $26 \pm 2$ | | Wang et al. 2022 (this study) |

<sup>a</sup> Abbreviations: GST = glutathione S-transferase; MT = magnetic tweezers; SNAP, CLIP = two self-labelling protein tags that respectively react with small molecules O<sup>6</sup>-benzylguanine and O<sup>6</sup>-benzylcytosine; PBS = phosphate buffer saline; FH1 = an intrinsically disordered peptide derived from the FH1 domain of formin mDia1; I27 = the immunoglobulin-like domain of titin; AFS = acoustic force spectroscopy.
<sup>b</sup> Only the buffering system, the major salts, and the rapamycin concentration are provided. Among the other components one may find DTT and EDTA (at millimolar concentrations) and/or organic solvents used to dissolve rapamycin (e.g. methanol, ethanol, or DMSO at up to a few % v/v).
<sup>c</sup> The dissociation equilibrium constants for the FKBP12•rapamycin and FRB•rapamycin complexes have been reported between $\simeq 0.2$ and $\simeq 2$ nM and between 2 and 20 nM, respectively (see the values compiled in Kostrz et al. 2019 [1] and in Joshi et al. 2022 [21]). For rapamycin concentrations ranging from 10 to 100 nM, it is assumed that FKBP12 is nearly fully occupied by the macrolide whereas FRB is nearly free of it [14,16]: one studies the interaction between FKBP12•rapamycin and FRB.
<sup>d</sup> The temperature at which data were collected is sometimes not specified (NS). However, since we previously measured the activation energy for the dissociation reaction, it is possible to evaluate the variations of $k_{\text{off}}$ with $T$ using to Arrhenius law. With $E_{\text{off}} \simeq 60 \text{ kJ/mol}$ , between 10 and 30 °C and taking $k_{\text{off}} = 30 \text{ s}^{-1}$ at $T=21.7$ °C as an example [1], one would have $k_{\text{off}} = 26 \text{ s}^{-1}$ at $T=20$ °C and $39 \text{ s}^{-1}$ at $T=25$ °C.
<sup>e</sup> In this table $k_{\text{off}}$ identifies with $k_{\text{off}}^0$ , the dissociation rate constant at zero force obtained in force spectroscopy
measurements. <sup>f</sup> Data given for different replicates.
<sup>g</sup> Both FRB and FKBP12 are here attached to the J-DNA scaffold by their N-terminus. Extra data collected after engraftment of the proteins on internal amino acids are provided in Table S4: the obtained dissociation rate constants at zero force lays in the range of the values displayed in present Table S3.
<sup>h</sup> Inverse-variance weighted mean on four measurements achieved at 0.05, 0.1, 0.2, and 500 nM rapamycin.
<sup>i</sup> FRB and FKBP12 are attached to their handles by their N- and C-terminus, respectively.
<sup>j</sup> A biexponential fit yielded two characteristic dissociation times and thus two dissociation rate constants; by combination with the single association rate constant it led to two dissociation equilibrium constants. Only one concentration point, at 1 nM FKBP12, was used in this study.

**Table S4:**
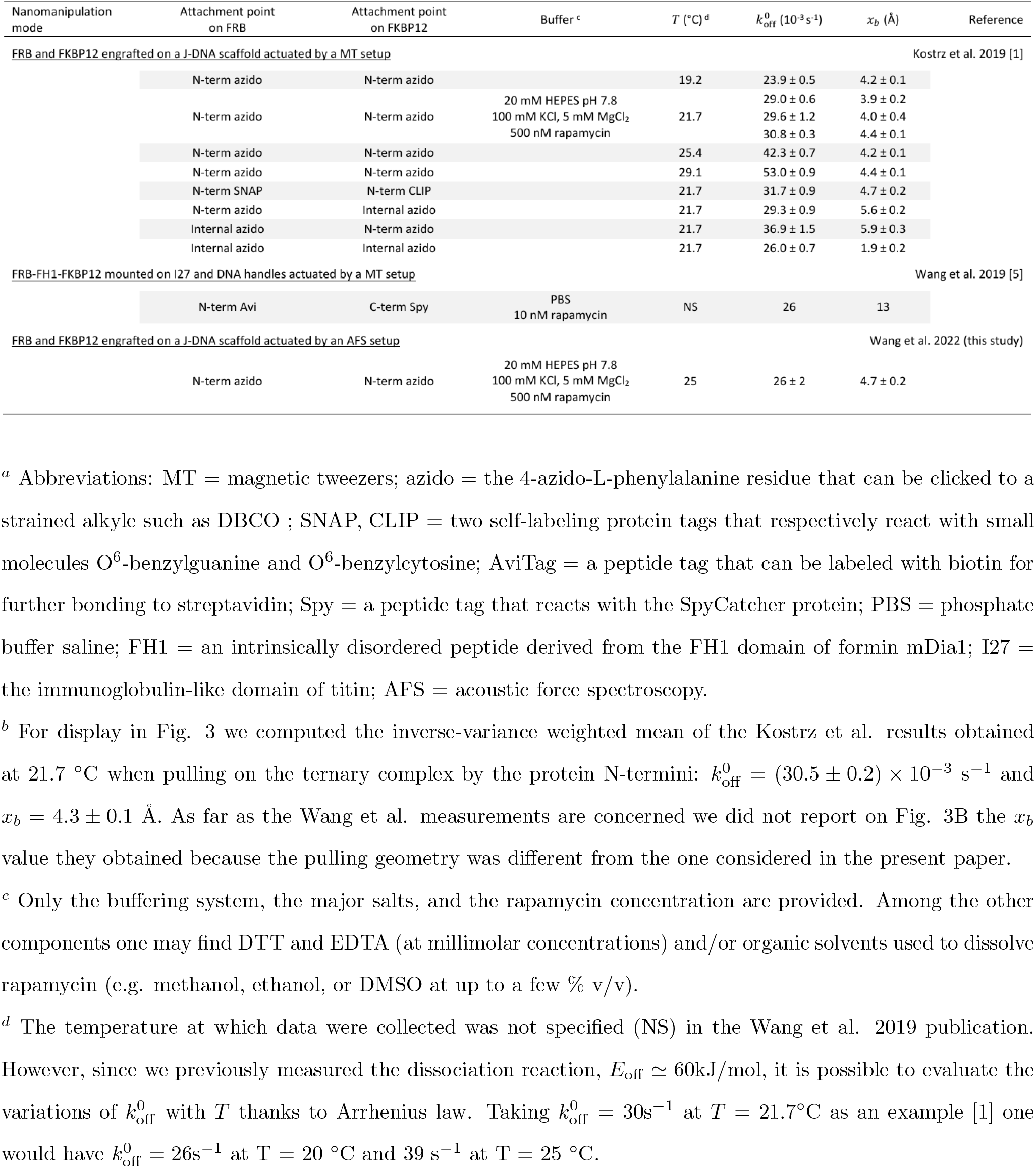
Published single-molecule force spectroscopy parameters for the FKBP12•rapamycin•FRB ⇌ FKBP12•rapamycin + FRB dissociation. 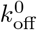 corresponds to the dissociation rate constant at zero-force and *x*_*b*_ to the distance from the potential well to the barrier of the energy landscape of interaction, both obtained from fitting data with the Bell model (Eq. 7) ^*a,b*^.

### Supplementary Figures

**Figure S1:**
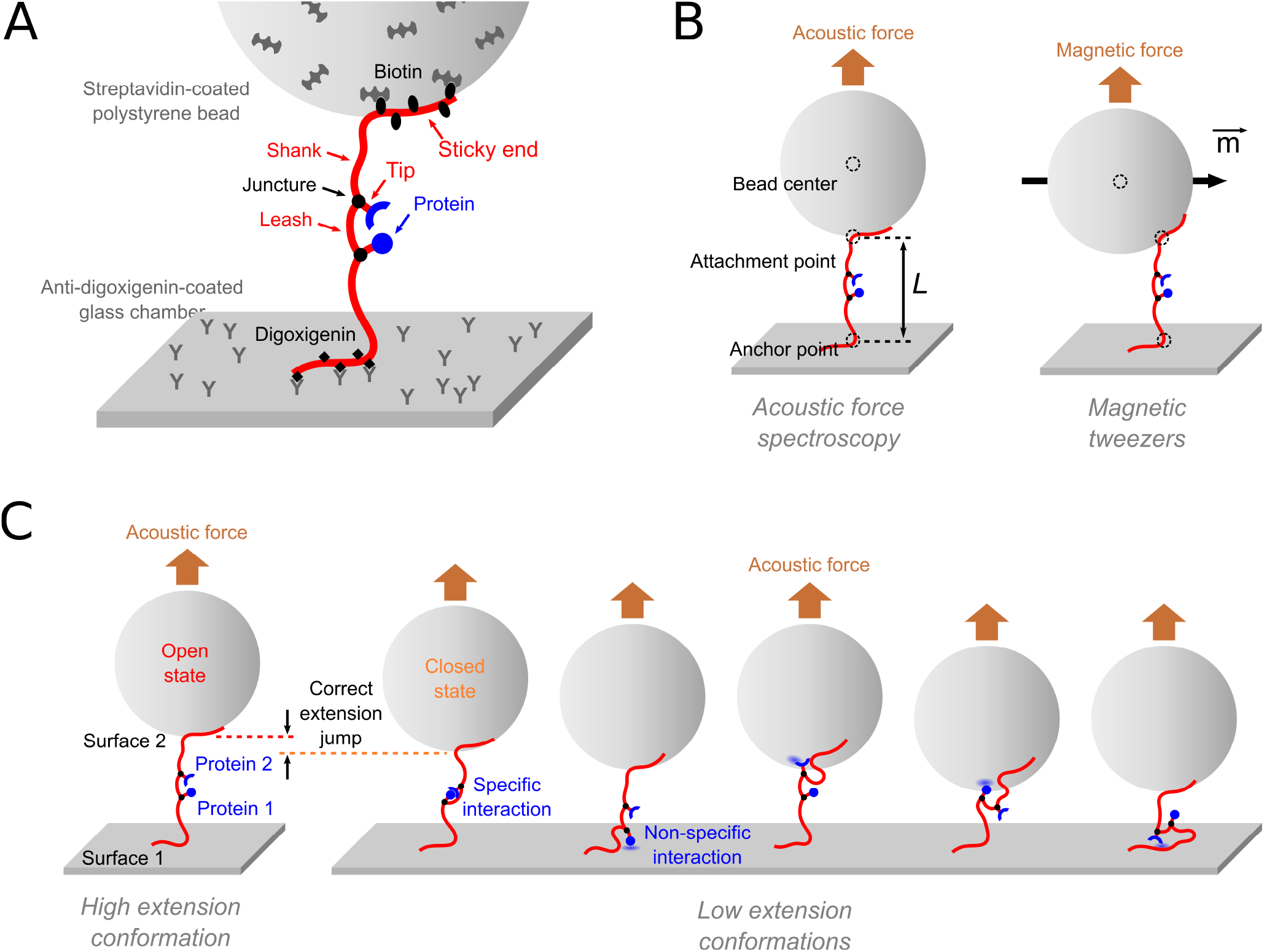
Diagrams of the J-DNA anatomy and working principles. (A) The scaffold is made of three continuous dsDNA helices: two branches and one leash that connected them at the junctures. The segments of the branches comprised between the junctures and the protein engraftment points are called the tips; the ones comprised between the junctures and the digoxigenin- or biotin-labelled sticky ends are called the shanks (see Table S1 for dimensions). To enable micromanipulation by acoustic force spectroscopy the J-DNA are attached to two handles, the surface of the anti-digoxigenin-coated chamber and a streptavidin-coated polystyrene bead (the scaling between the latter particle and the nucleic acids construct has been maintained on the drawing). (B) The bead being free to rotate in the acoustic force field, it is straightforward to measure the J-DNA extension *L*. Indeed, the attachment point to the bead always lays between the particle center and the anchor point at the chip surface. Conversely, in the case of magnetic tweezers, the particle moment must align with the field; such angular constraint results in an attachment point not necessarily located in between the bead center and the anchor point, which impedes the determination of tether length when *L* ≃ *R*, with *R* the bead radius. (C) The contour lengths of the various J-DNA segments were chosen to ensure a unique signature to the extension jumps associated with protein-protein bond ruptures. Indeed, after disruption of an interaction between one of the proteins and one of the surfaces one should observe a much larger step variation than expected (see Table S2 for numerical values).

**Figure S2:**
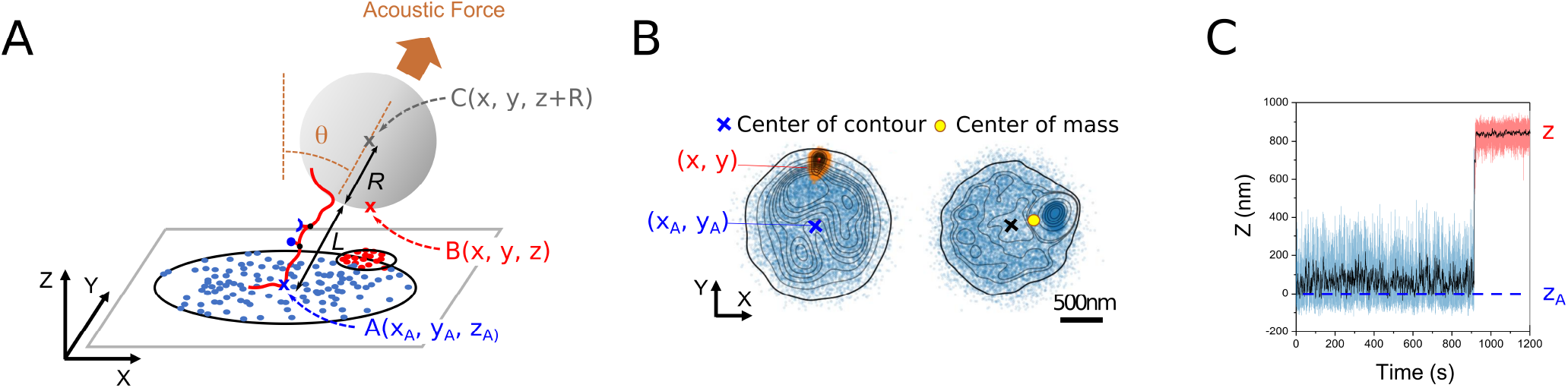
(A) Geometrical notations to determine anchor point A, scaffold length *L* and pulling angle *θ* (see Eqs. 1 and 2). The blue points are the projection of bead position on the (x,y) plane when low or zero force is applied. The red points are the projection of bead positions when an acoustic force is applied. (B) Left: Illustration of the effect of pulling angle *θ* on the lateral shift between anchor point A and bead position B at high force. Right: Illustration of the use of a density kernel on lateral positions to determine the anchor point as center of the external density map contour, in case the trace exhibits spurious sticking points. (C) Graphical representation of the convention to establish the zero altitude (dashed line) after time average of z trace fluctuations (in black).

**Figure S3:**
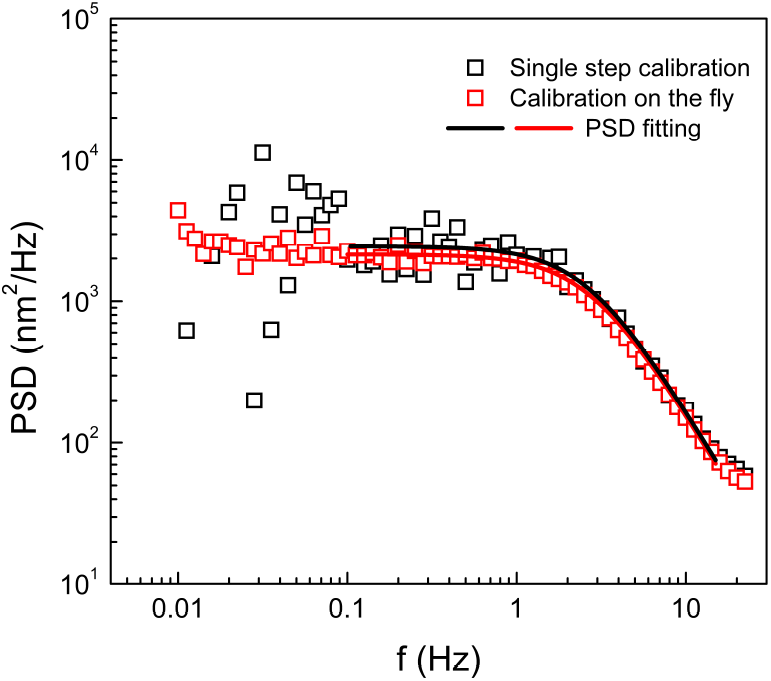
Comparison of power spectra densities for single step vs on the fly calibration. Example of ID1 for the FKBP12 and FRB proteins.

**Figure S4:**
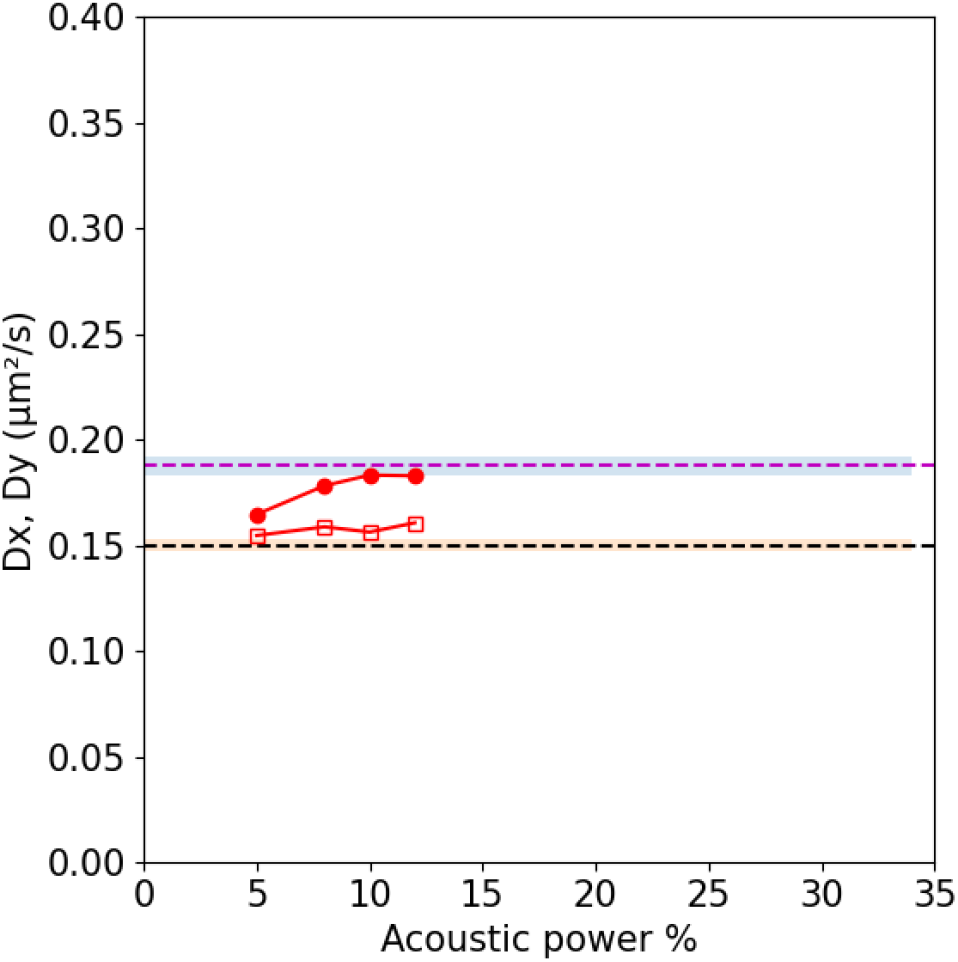
Diffusion coefficient from PSD fit as a function of tested power for a bead tethered by J-DNA functionalized with the FKBP12 and FRB proteins. Circles: individual PSD^*x*^ fit with one *D* and one *k*, when the condition *f*_*c*_ *< f*_*max*_*/*2 is fulfilled. Squares: individual PSD^*y*^ fit. Horizontal lines: diffusion coefficient calculated from the unique value of *R*_*x*_, *R*_*y*_ for each bead, as determined by the global fit of PSD. The shaded stripe indicates the error bar, based on the error on the fitting value of *R* and the dependence of *D* on J-DNA extension at each power.

**Figure S5:**
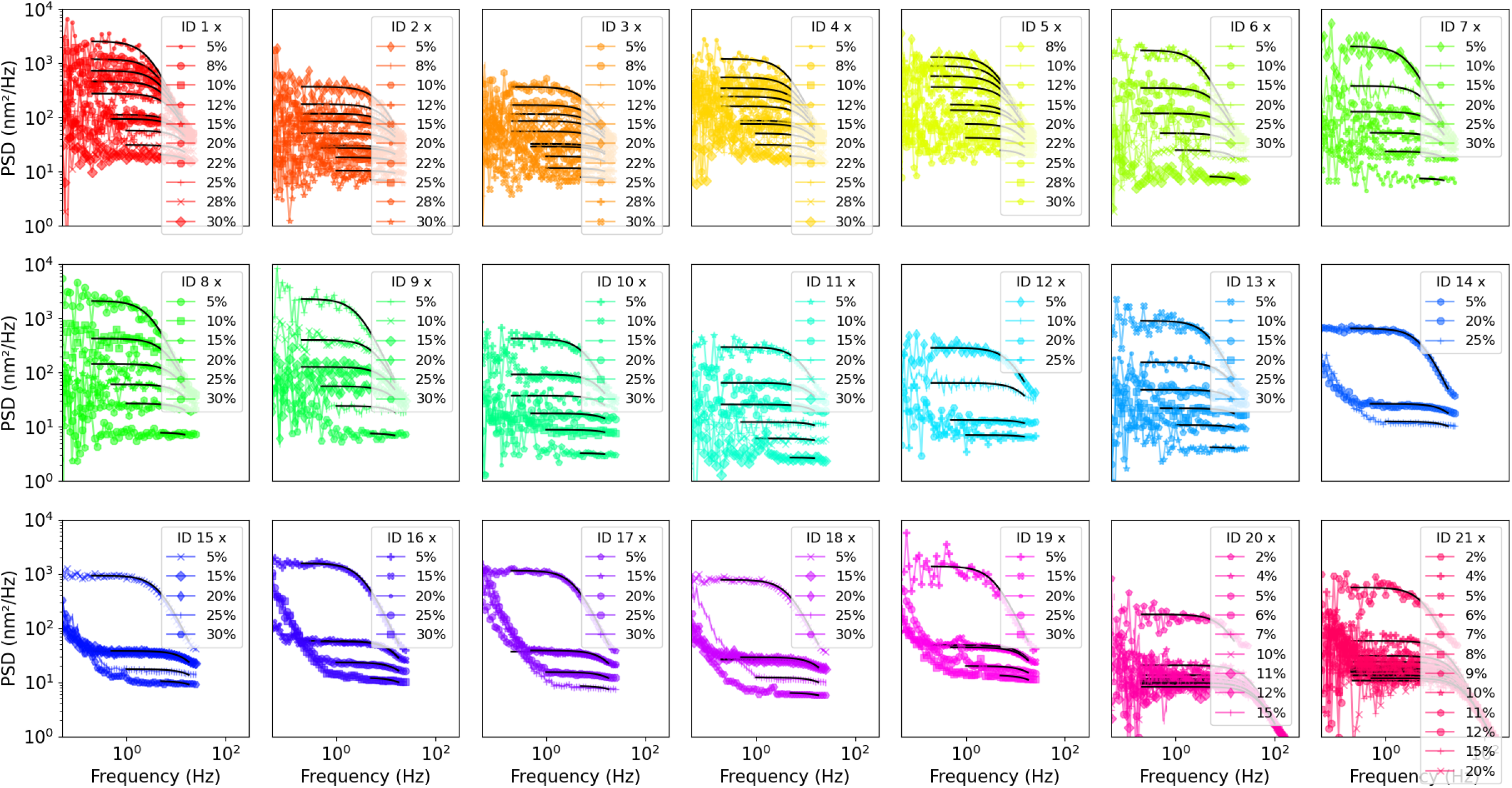
Power spectra density of *x* fluctuations for 21 beads tethered by J-DNA functionalized with the FKBP12 and FRB proteins and subjected to different acoustic powers in % units. Fits to Eq. 3 and 4 are superimposed, taking for each bead as fitting parameters : the bead radius *R*^*x*^ and the pendulum stiffness 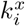 for each power i. The fitting range is chosen as to contain the plateaus of all curves as well as the corner frequency *f*_*c*_, if the condition *f*_*c*_ *< f*_*max*_*/*2 is fulfilled.

**Figure S6:**
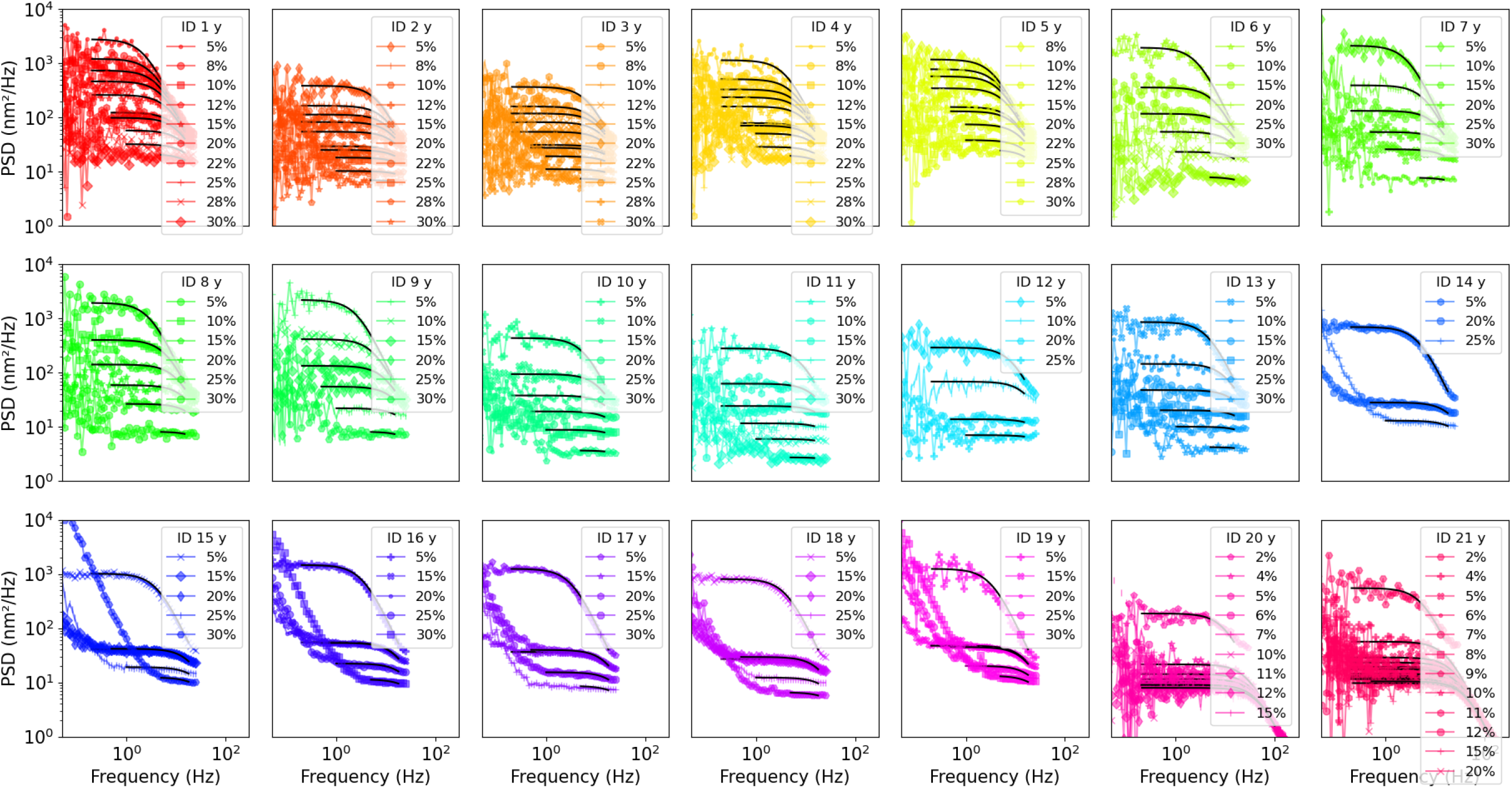
Power spectra density of *y* fluctuations for 21 beads tethered by J-DNA functionalized with the FKBP12 and FRB proteins and subjected to different acoustic powers in % units. Fits to Eq. 3 and 4 are superimposed, taking for each bead as fitting parameters : the bead radius *R*^*y*^ and the pendulum stiffness 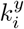 for each power i.

**Figure S7:**
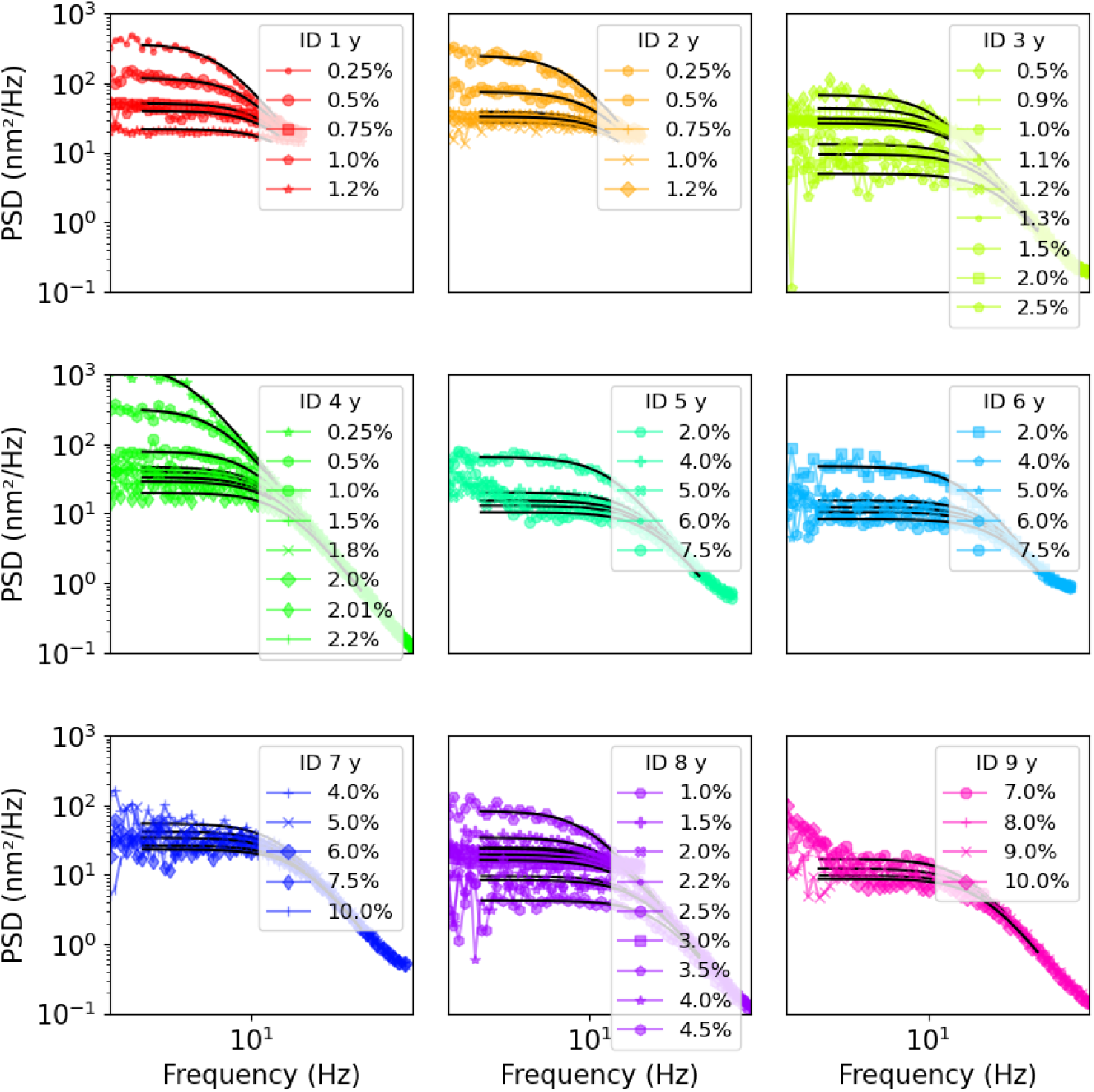
Power spectra density of *y* fluctuations for 9 beads tethered by J-DNA functionalized with the Nef and Nef19 protein and subjected to different acoustic powers. Fits to Eq. 3 and 4 are superimposed, taking for each bead as fitting parameters : the bead radius *R*^*y*^ and the pendulum stiffness 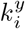 for each power i.

**Figure S8:**
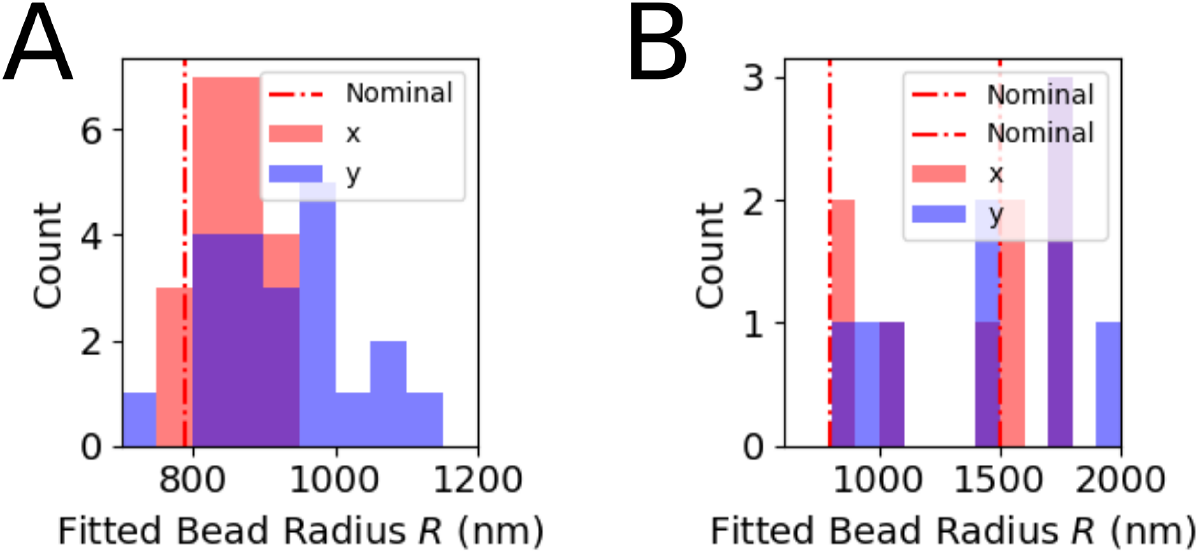
Distribution of apparent bead radius (in nm) retrieved from global PSD fit. The nominal bead radius is indicated by the red dashed line. (A) 21 beads tethered by J-DNA functionalized with the FKBP12-rapamycin-FRB complex. (B) 9 beads tethered by J-DNA functionalized with the Nef-Nef19. Two bead sizes were used in these measurements, represented by 2 red dashed lines, and matching with fitted radii.

**Figure S9:**
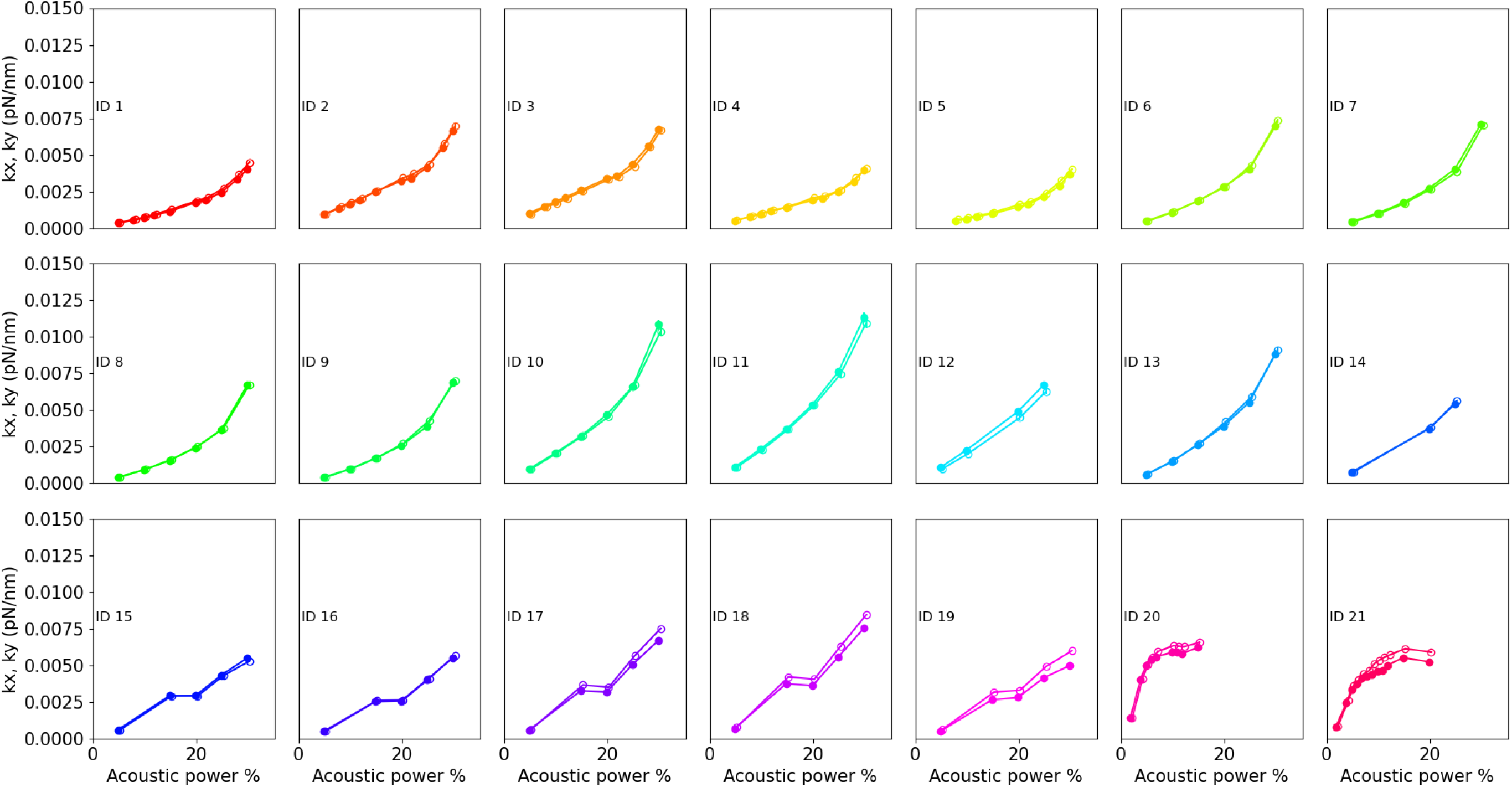
Stiffness of the pendulum from global PSD fit as a function of acoustic power for 21 beads tethered by J-DNA functionalized with the FKBP12 and FRB proteins. Plain symbols: stiffness obtained from PSD of *x* fluctuations. Empty symbols: stiffness obtained from PSD of *y* fluctuations.

**Figure S10:**
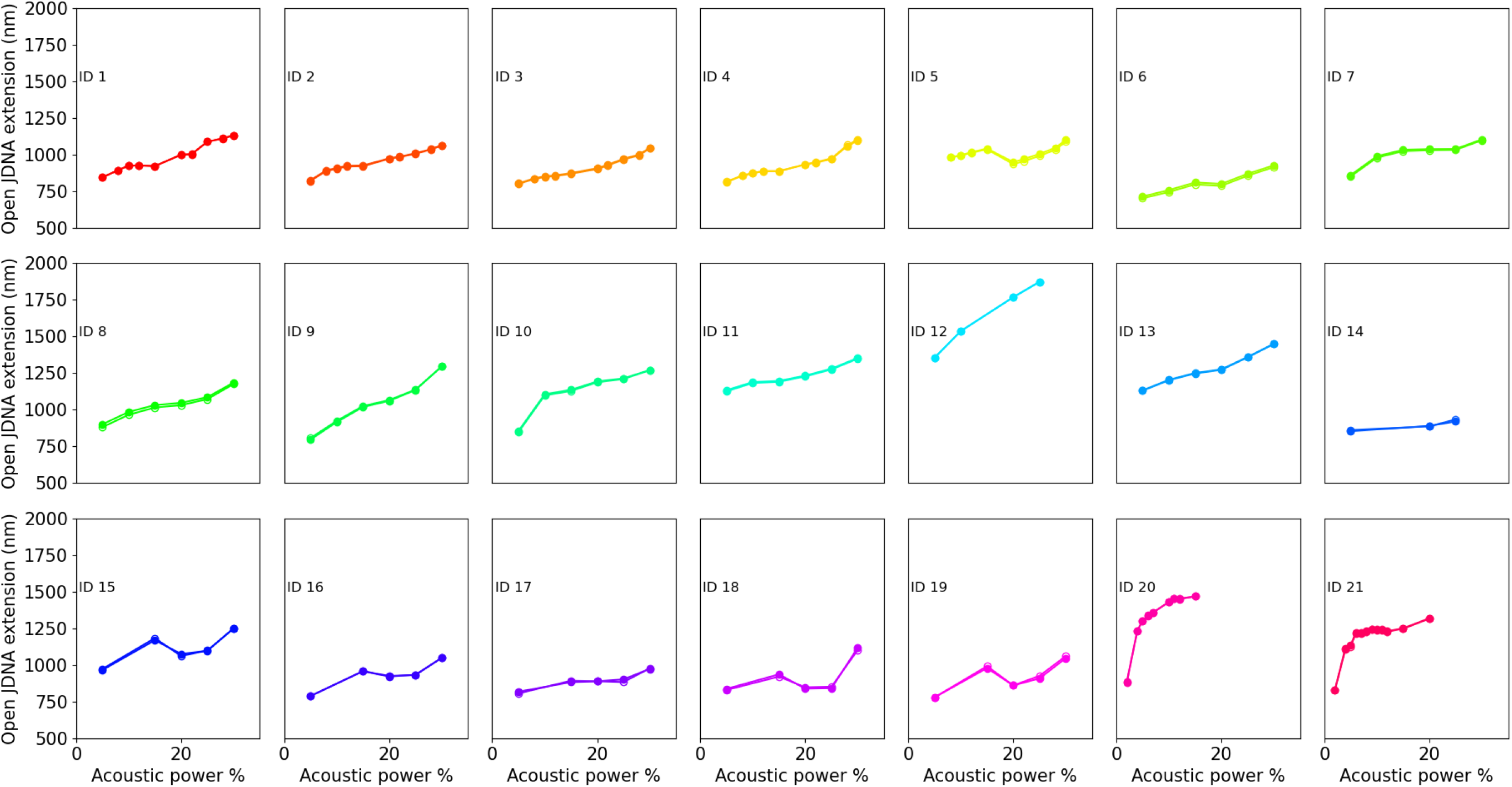
Open J-DNA extension (in nm) as a function of acoustic power for 21 beads tethered by J-DNA functionalized with the FKBP12 and FRB proteins and subjected to different acoustic powers.

**Figure S11:**
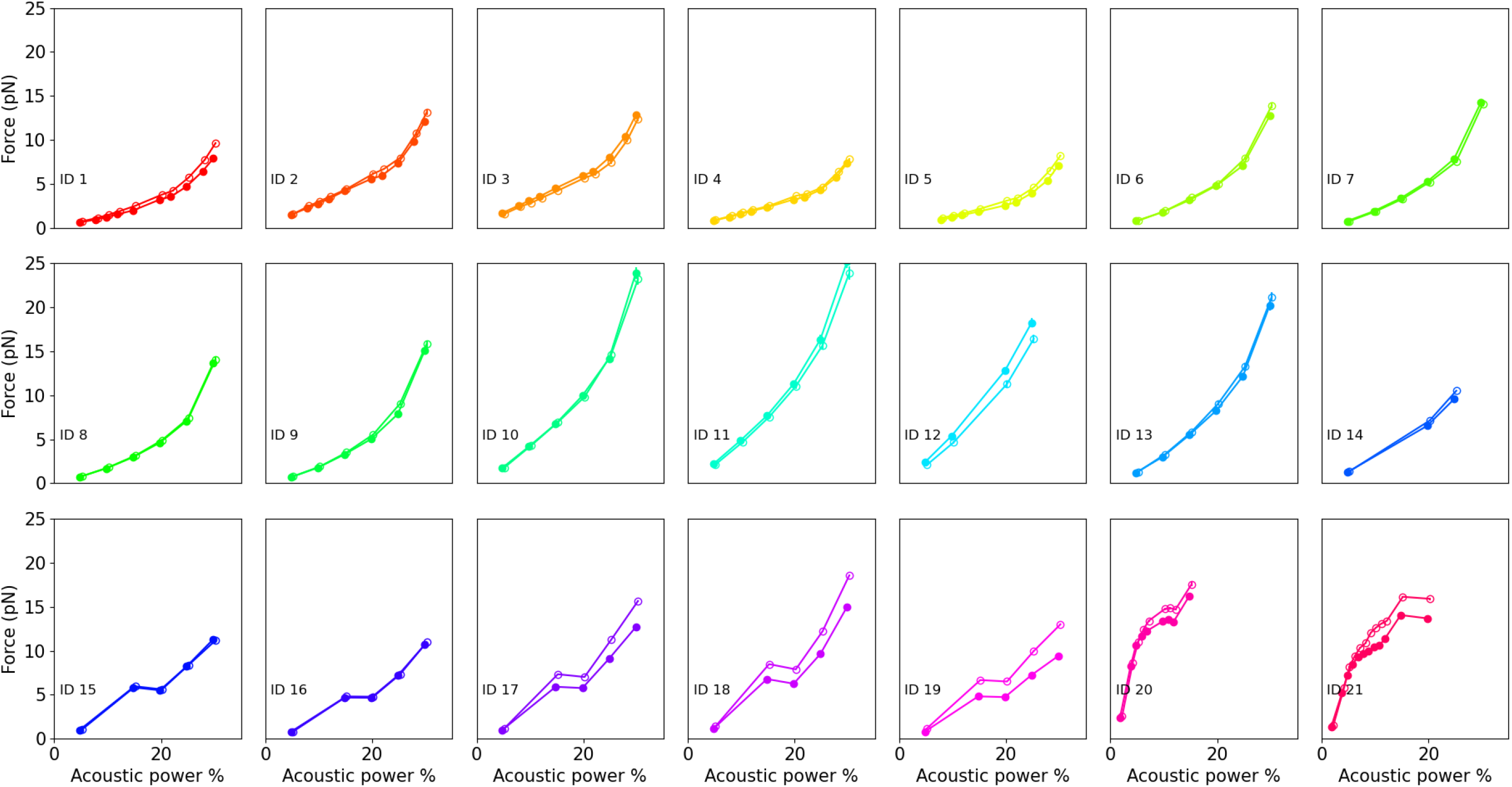
Force (in pN) as a function of acoustic power for 21 beads tethered by J-DNA functionalized with the FKBP12 and FRB proteins.

**Figure S12:**
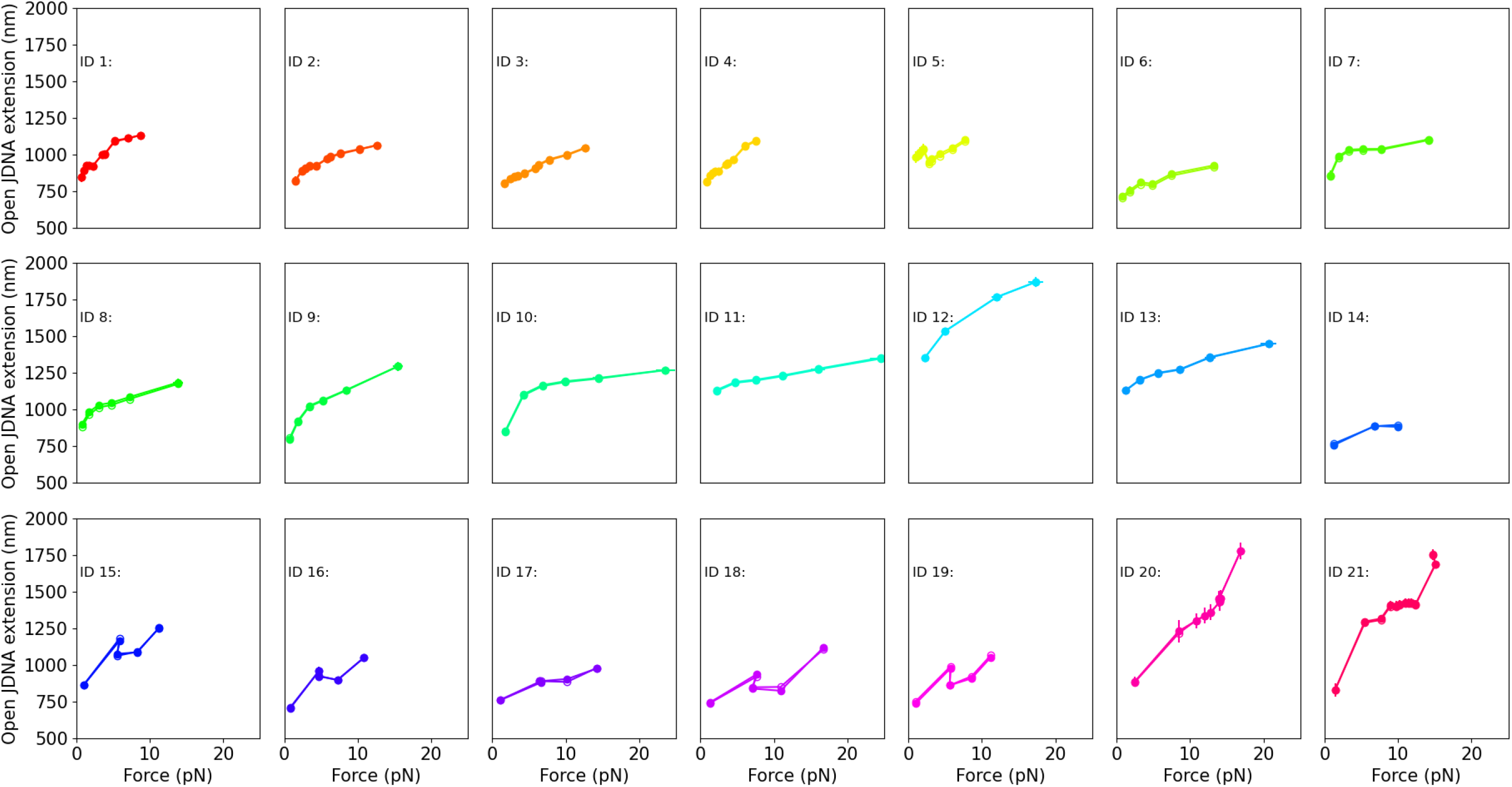
Open J-DNA extension (in nm) as a function of force for 21 beads tethered by J-DNA functionalized with the FKBP12 and FRB proteins and subjected to different acoustic powers.

**Figure S13:**
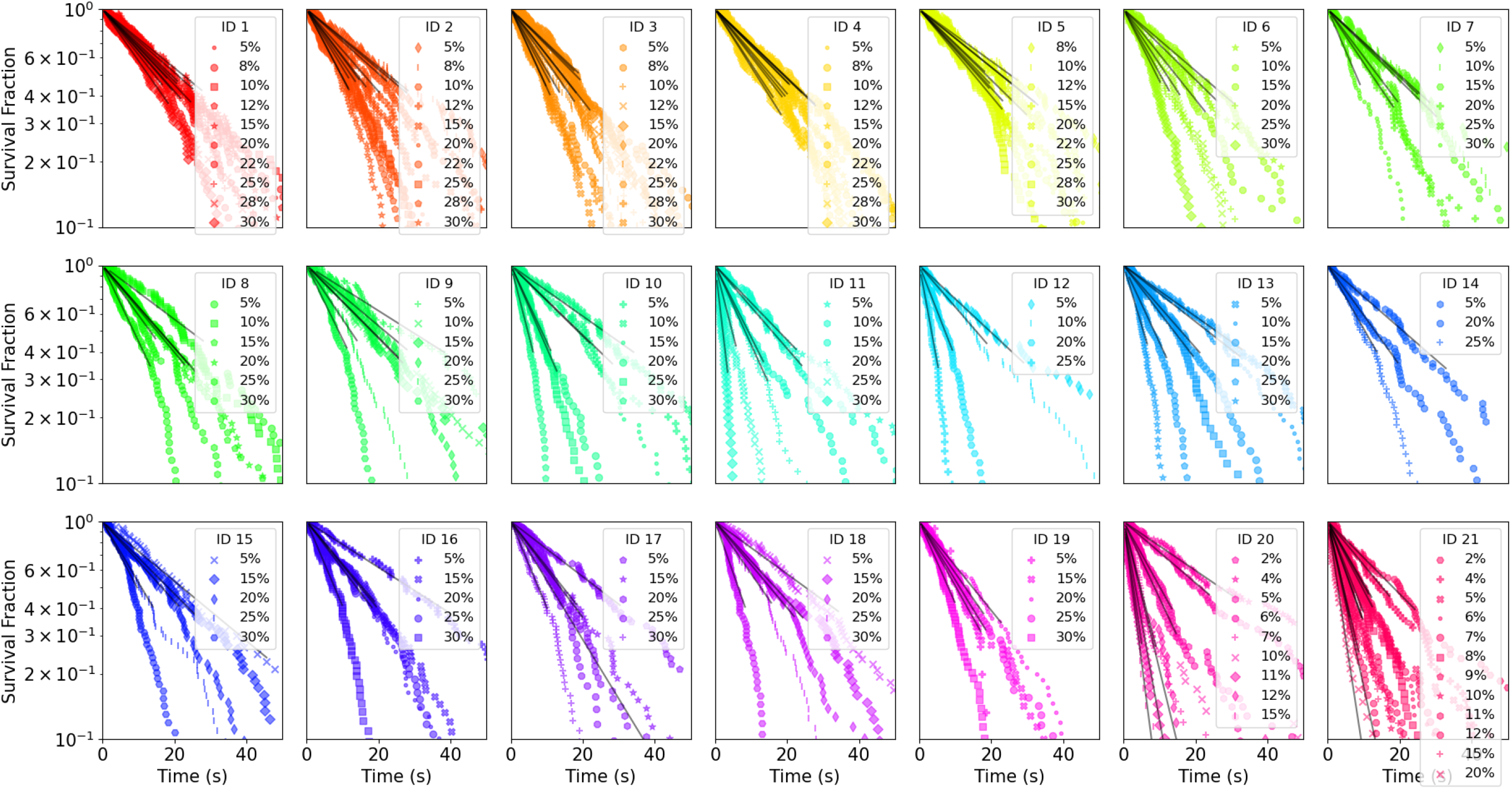
Survival of FKBP12-rapamycin-FRB bonds for 21 complexes and various acoustic power (in %). Monoexponential fits are superimposed.

**Figure S14:**
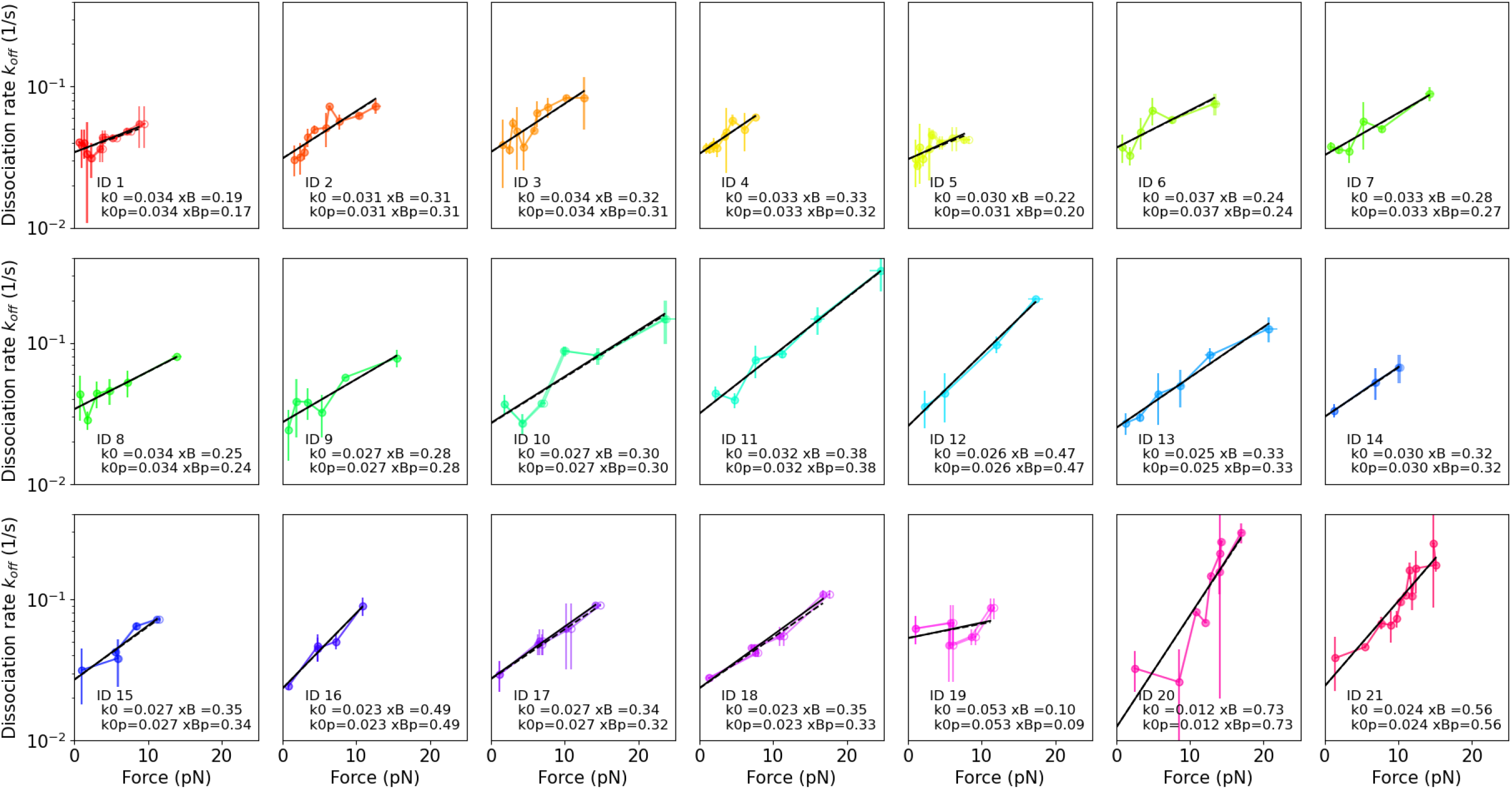
Dependence of off-rate of FKBP12-rapamycin-FRB bonds as a function of force for 21 individual complexes (plain symbols). Plain line indicates fit with the Bell model (Eq. 7). Empty symbols are calculated using the force projected along the main axis of the scaffold, considering the angle *θ*. The dashed line represent the Bell fit for the projected force. Bell parameters 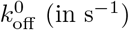 and *x*_*b*_ (in nm) are indicated in each case for non-projected and projected force.

**Figure S15:**
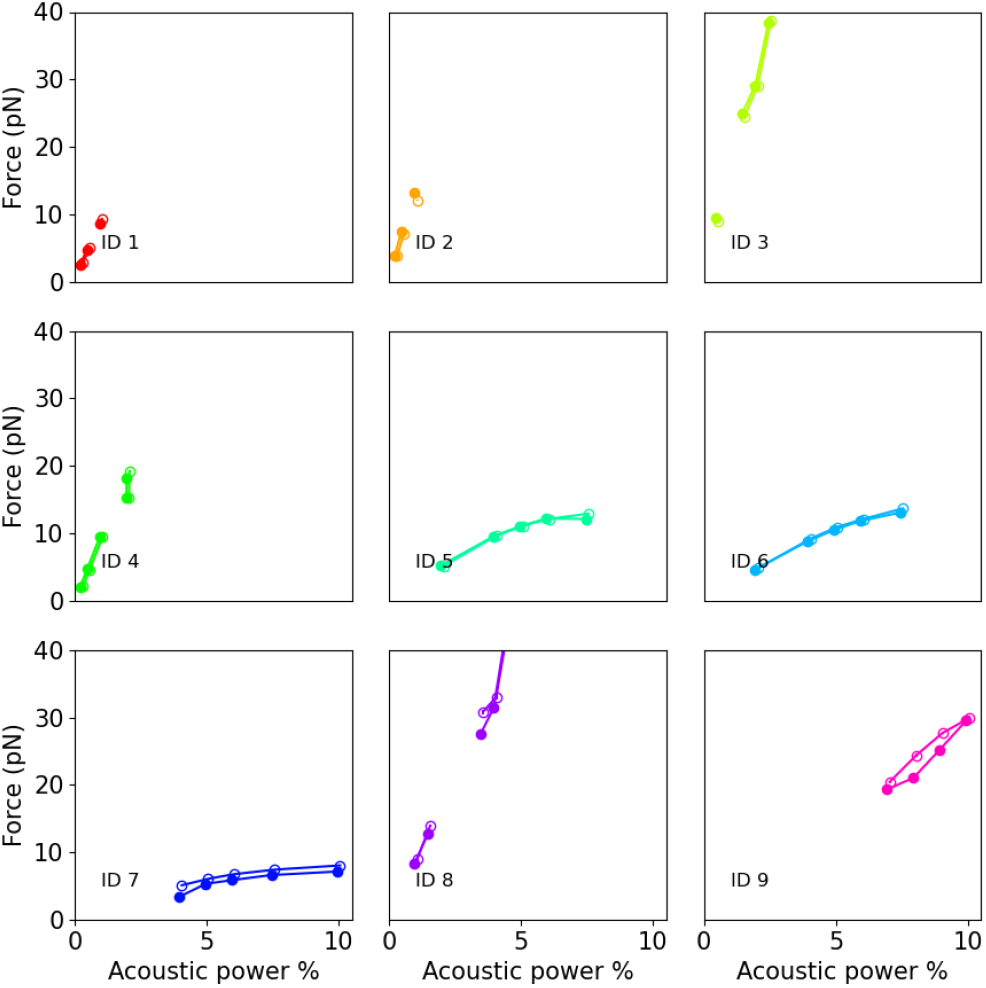
Acoustic force (in pN) as a function of test power for 9 beads tethered by J-DNA functionalized with the Nef-Nef19 complex. Plain symbols: force obtained by PSD in *x* direction. Empty symbols: force obtained by PSD in *y* direction. Notice that ID 5,6,7 correspond to *R* = 900 nm beads, while other IDs correspond to *R* = 1500 nm.

**Figure S16:**
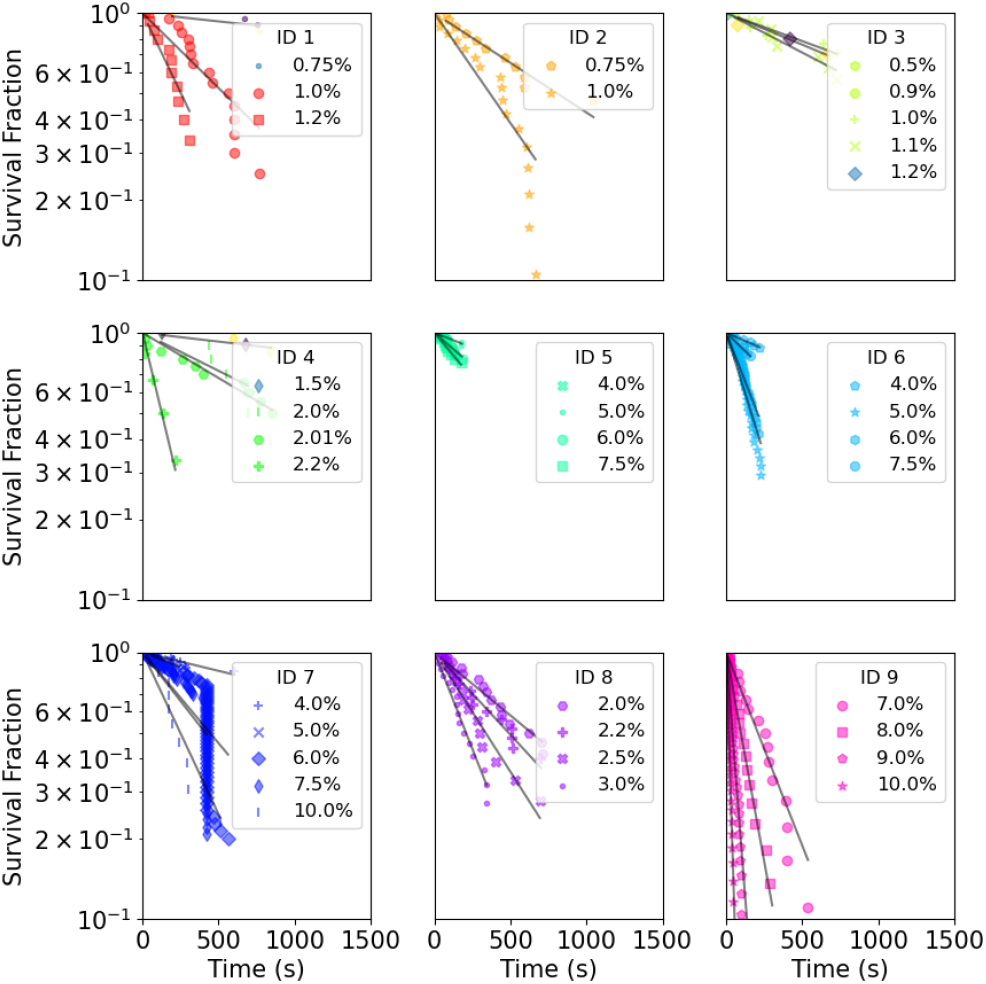
Survival of Nef-Nef19 bonds for 9 complexes and various acoustic power (in %). Vertically aligned points represent durations which have been censored by the measurement duration. Monoexponential fits are performed on the time range limited by the measurement duration and are superimposed.

**Figure S17:**
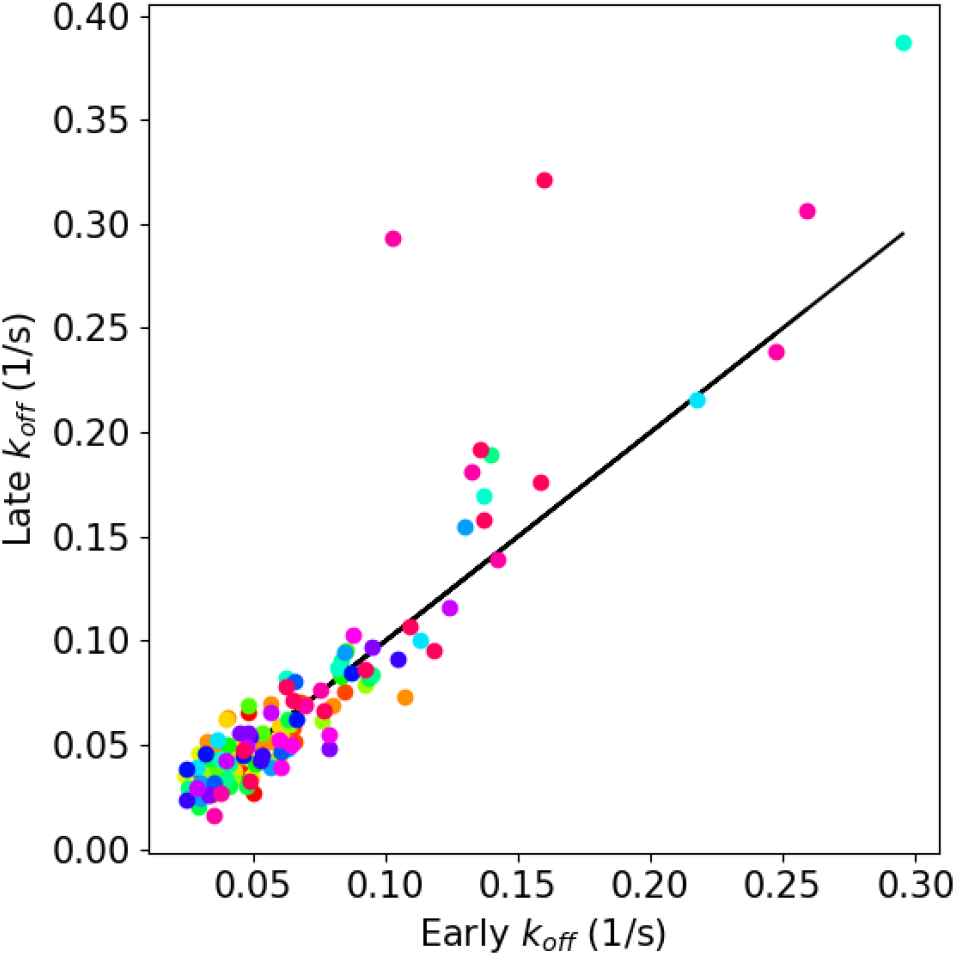
Unbinding kinetics for the FKBP12-rapamycin-FRB complex is not affected by repetitive force cycling and the succession of association and dissociation events. The off-rate is calculated for each of the 21 molecules and each applied acoustic power based on the lifetime of the 50 first unbinding events (early *k*_off_) or 50 last unbinding events (late *k*_off_). The black line represents *x* = *y*.

